# MERTK and RAC1^P29S^ Drive Melanoma Cannibalism of T cells

**DOI:** 10.64898/2026.11.11.766025

**Authors:** Lauren Penfield, Maddalena Nano, Denise Montell

## Abstract

Metastatic melanoma cells are cannibalistic, engulfing living and dead lymphocytes, but the molecular mechanisms that drive this behavior, including how they recognize their targets, are unknown. RAC is a Rho family GTPase that plays a key role in F-actin dynamics. RAC1^P29S^ is a hyperactivating driver mutation that is common in melanoma where it promotes proliferation, motility, and therapy-resistance. Here we show that RAC1^P29S^ melanoma cells engulf more Jurkat T cells than an isogenic RAC1^WT^ revertant line. Live imaging captured lamellipodial dynamics and whole-cell engulfment in real time. An inhibitor screen across RAC1^P29S^ and RAC1^WT^ patient-derived lines showed that cannibalism requires the phosphatidylserine receptor MERTK. A RAC1^P29S^ line lacking detectable MERTK was not cannibalistic, indicating that RAC1 hyperactivity amplifies a specific, receptor-mediated response. RAC1^P29S^ lines are sensitive to RAC and Arp2/3 inhibition. We propose a “grip and gulp” model in which MERTK provides the grip and Arp2/3-driven branched actin the gulp.

## Introduction

Melanoma is the 5th most common cancer in the United States and is the deadliest type of skin cancer. Whereas melanomas detected early have good prognosis, the five-year survival for metastatic melanoma is 32% ^1^. Understanding the properties of metastatic melanomas and how to effectively target them will be critical to improve treatment options.

An understudied yet striking feature of metastatic melanoma cells is that they display cellular cannibalism, defined as one cell eating another, whole or in part, living or dead, whereas melanocytes and primary melanoma cells do not phagocytose other cells ^2,3^. Lugini et al reported cell-in-cell structures in tissue sections of metastatic melanoma tumor samples, indicating cells engulfing other cells ^2^. In vivo, T cells can be found inside of melanoma cells, suggesting that melanoma cells cannibalize T cells. In culture, metastatic melanoma cells engulf apoptotic bodies at similar rates as macrophages ^4^, and they can phagocytose living lymphocytes ^2^. Over 25 different types of cancers, including breast and pancreatic ductal adenocarcinoma, exhibit cell-in-cell structures ^3–6^. Engulfment of other cells may facilitate tumor cell survival both by supplying nutrients in depleted tumor microenvironments and suppressing anti-tumor immune responses locally.

The molecular mechanisms that control melanoma cell cannibalism remain to be fully elucidated. Ezrin, caveolin, TM9SF4, and actin have been implicated ^2,4,7^; but how these proteins contribute to engulfment and whether additional signaling pathways are required remain unclear. Most importantly, how melanoma cells recognize the cells they engulf is unknown. Because melanoma cells internalize inert latex beads, the prevailing view is that cancer cell cannibalism proceeds non-specifically, without target recognition at all. RAS-related C3 botulinum toxin substrate 1 (RAC1) is a small

GTPase that functions as a key regulator of actin polymerization and protrusion, cell migration, and phagocytosis, and RAC1 activity is often elevated in cancer ^8–11^. Overexpression of RAC1 itself or of RAC1 activators, such as Guanine nucleotide Exchange Factors (GEFs) increases RAC1 activity. Mutation or reduced expression of GTPase-activating proteins (GAPs) that inactivate RAC1 can achieve a similar effect ^12^. Mutations that directly activate RAC1 are prevalent in a subset of cancers ^12,13^, and the activating RAC1^P29S^ mutation is common in metastatic melanoma; only activating mutations in BRAF and NRAS are more frequent ^14^. The UV-induced, RAC1^P29S^ mutation induces spontaneous RAC1 activation and fast GDP-GTP cycling ^15^. RAC1^P29S^ increases proliferation, migration, and therapy-resistance of melanocytes ^14^. However, the effects of RAC1^P29S^ on melanoma cell engulfment behaviors, if any, have not been reported.

Rac hyperactivity can amplify cell-eating behavior in other contexts. In the Drosophila ovary, for example, constitutively active Rac1^G12V^ causes border cells to engulf neighboring polar cells whole ^16^, whereas wildtype Rac drives F-actin protrusion and border cell migration ^11^. In this context, Rac1^G12V^ converts normal, sublethal nibbling of substrate nurse cells to lethal phagocytosis ^16,17^. Both sublethal and lethal engulfment of nurse cells require the receptor Draper, the fly homolog of the mammalian engulfment receptor MEGF10 ^16,17^. In non-phagocytic HeLa cells engineered to express a phosphatidylserine (PS)-binding domain and a photo-activatable Rac GEF, illumination drives engulfment of PS-positive apoptotic bodies ^18^. Hyperactivation of Rac2 also causes engineered chimeric-antigen-receptor-expressing macrophages to engulf more whole target cells ^16^. Whether cancer cells might be able to naturally evolve a similar combination of an activating Rac mutation and receptor expression, and thereby acquire the capacity to consume other cells, was not known.

Here we identify MERTK as an engulfment receptor required for melanoma cell cannibalism, and show that an endogenous, patient-derived, oncogenic RAC1^P29S^ mutation enhances this behavior in human tumor cells. We report that in the presence of MERTK, a receptor for PS, RAC1^P29S^ enhances melanoma cell phagocytosis of dead and living T cells. MERTK overexpression is associated with enhanced cannibalism even in RAC1^WT^ melanoma cells. Pharmacological inhibition of MERTK, F-actin, and PI3 kinases also broadly reduced cannibalism, while RAC1^P29S^ lines were more sensitive to RAC and Arp2/3 inhibition than RAC1^WT^ lines. Cannibalism in melanoma cells is also sensitive to inhibition of RAS-signaling pathways. Inhibitors of myosin, PAK or AKT enhanced cannibalism in a subset of lines. Overall, our work shows that melanomas harness MERTK and Rac-mediated F-actin polymerization to enhance cell cannibalism.

## Results

### The RAC1^P29S^ mutation promotes melanoma cell engulfment of dead T cells

To determine the role of *RAC1*^*P29S*^ in cell cannibalism, we used an isogenic pair of melanoma lines. The patient-derived line IGR1 has an endogenous *RAC1*^*P29S*^ mutation and a BRAF^V600E^ mutation (IGR1-RAC1^P29S^). In the line called IGR1-RAC1^WT^, *RAC1*^*P29S*^ has been reverted to *RAC1*^*WT 19*^. Melanoma cells were labeled with Celltrace-CFSE and plated (Fig. 1A, green). Then heat-killed (56°C, 30 min) Jurkat T cells, a leukemia T cell line, were labelled with Celltrace-FarRed and added at a ratio of 5:1 (target:melanoma) (Fig. 1A, A’ magenta). Heat killing produces primary necrosis and has been used previously as a defined model of unscheduled necrotic death ^20^. Necrosis is common in solid tumors and is present in approximately one quarter of nodular cutaneous melanomas, where it is associated with greater tumor thickness and reduced survival ^21^. We mixed melanoma and T cells for 24 hours, fixed the samples, and measured the percentage of melanoma cells that engulfed at least one T cell to compare the original IGR1-RAC1^P29S^ cells (Fig. 1B) and the reverted IGR1-RAC1^WT^ cells (Fig. 1C). About 30% of IGR1-RAC1^P29S^ cells engulfed heat-killed Jurkat T cells (Fig. 1D), with evidence of individual melanoma cells engulfing multiple targets (Fig. 1B). Significantly fewer IGR1-RAC1^WT^ cells engulfed any target (Fig. 1C-D), and phagocytosis of multiple targets was not observed.

**Figure 1.**
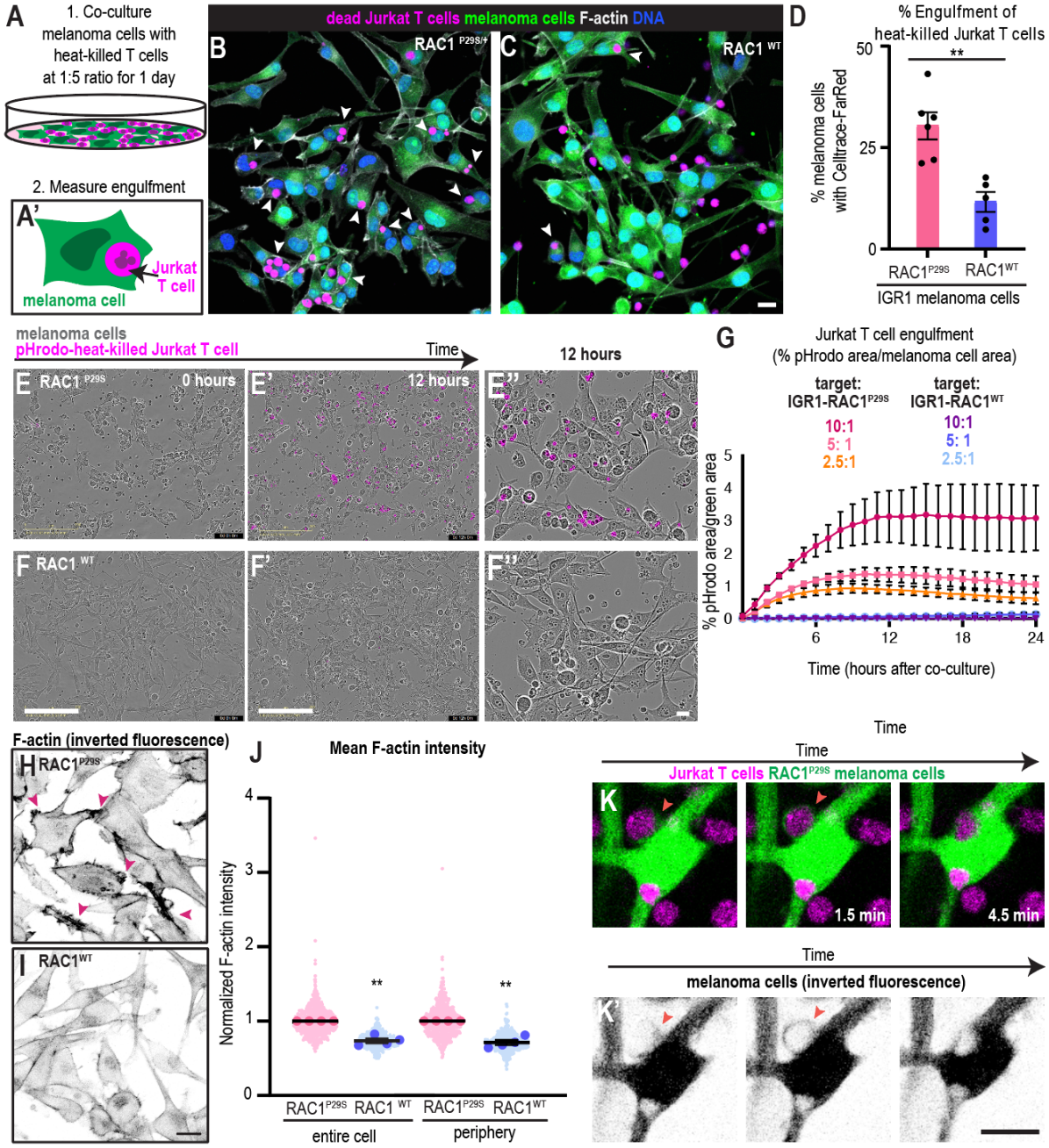
The RAC1^P29S^ mutation enhances melanoma cell engulfment of necrotic T cells. A. Schematic of experimental setup for melanoma cell-T cell co-cultures. B-C Confocal images of melanoma cells labeled with CellTrace-CFSE(green) incubated with heat-killed Jurkat T cells labeled with Celltrace-Far Red (magenta). Cells were stained with Hoechst (blue) to label nuclei and Phallodin to label (F-actin). B shows IGR1 cells with the endogenous RAC1^P29S^ mutation. C shows IGR1 revertant line with RAC1^WT^. Scale bar: 20 microns. D. Plot showing the mean (column) +/-standard error of the mean (SEM, black bars) percentage of melanoma cells that engulfed a heat-killed Jurkat as measured by the presence of CelltraceFar Red inside the melanoma cell for the indicated conditions. Each dot represents an experimental replicate. N=6 experimental replicates for RAC1^P29S^ and N=5 for RAC1^WT^. A student’s T test was performed, **=p<0.01. E-F Images from timelapse imaging co-cultures of IGR1-RAC1^P29S^ (E) or IGR1-RAC1^WT^ melanoma cells and heat-killed Jurkat T cells labeled with pHrodo that fluoresces when acidified (magenta) at time 0 (E-F) and 12 hours (E’-F’). E’’-F’’ are magnified images at the 12 hour timepoint. G. Plot showing the mean % pHrodo positive area over the green area for the indicated conditions. N=4 experimental replicates for each condition. H-I.Confocal images showing the inverted fluorescence F-actin(Phalloidin) for H. IGR1-RAC1^P29S^ and I. IGR1-RAC1^WT^. Scale bar: 20 microns. J. Plot showing the normalized total mean intensity +/-SEM (black bars) and individual experimental means (large dots), and individual cell means (small dots). Each experimental replicate was normalized to the mean fluorescent intensity of IGR1P29S. A one sample T test was performed on experimental replicates. **=p<0.01. K. Images from a time lapse series of IGR1-RAC1^P29S^ cells labeled with CellTrace-CFSE co-cultured with Jurkat T cells labeled withCellTrace-FarRed (magenta). K’ shows the inverted fluorescence of CellTrace-CFSE, and orange arrows indicate membrane extensions that wrap around Jurkat T cells. Time is relative to the first image. Scale bar: 20 microns.

To investigate the effects over time, we performed imaging for 24 hours. We labeled heat-killed Jurkat T targets with pHrodo, a dye that fluoresces when internalized targets become acidified in the lysosome and is a readout for phagocytosis ^22^. Then, we added them to IGR1-RAC1^P29S^ and IGR1-RAC1^WT^ melanoma cells and imaged them every hour for 24-hours (Fig. 1E-F” and Supplementary Movies 1 and 2). In IGR1-RAC1^P29S^ cells, pHrodo accumulated over time (Fig 1E-E”) whereas little pHrodo accumulated in IGR1-RAC1^WT^ cells (Fig. 1F-F”).

We also tested a variety of target:melanoma cell ratios. At all target:melanoma ratios, IGR1-RAC1^P29S^ cells engulfed significantly more targets than IGR1-RAC1^WT^ (Fig. 1G and Fig. S1A and B, p<0.001). IGR1-RAC1^WT^ cells engulfed virtually no target cells at 2.5:1 and 5:1. At 10:1, IGR1-RAC1^WT^ engulfed a few targets for about 6 hours. In contrast, at lower target:melanoma ratios of 2.5:1 and 5:1, the IGR1-RAC1^P29S^ pHrodo signal plateaued at ∼12 hours (Fig. 1G). Increasing the target:melanoma ratio to 10:1 delayed the plateau specifically in IGR1-RAC1^P29S^ (Fig. 1G) indicating that the plateau was due to reduced target availability rather than saturated capacity for engulfment. We conclude that IGR1-RAC1^P29S^ cells engulf more heat-killed Jurkats and consume over a more prolonged period than IGR1-RAC1^WT^.

### Elevated cortical F-actin and phagocytic cup extension in RAC1^P29S^cells

We next investigated how IGR1-RAC1^P29S^ cells engulf their targets. Activated RAC1 binds the WAVE regulatory complex to stimulate Arp2/3-mediated branched F-actin polymerization ^23–25^. So, we compared F-actin levels and localization within IGR1-RAC1^P29S^ and IGR1-RAC1^WT^ cells. In RAC1^P29S^ cells, lamellipodia-like structures that were enriched in F-actin were prevalent (Fig. 1H, arrows) whereas F-actin was less intense in RAC1^WT^ cells (Fig. 1I). We quantified both peripheral and whole cell F-actin and found that both were increased in RAC1^P29S^ compared to RAC1^WT^ (Fig 1J). We conclude that RAC1^P29S^ promotes F-actin polymerization.

To observe membrane dynamics during engulfment, we performed live confocal imaging. Electron microscopy suggested that T cells sink into and are entrapped by melanoma cells rather than by extension of a typical phagocytic cup ^2^. However, phagocytic cups may be sensitive to fixation and the dynamics of the process had not been captured. Celltrace CFSE-labeled IGR1-RAC1^P29S^ or IGR1-RAC1^WT^ melanoma cells were co-cultured with Celltrace-FarRed-labeled Jurkat T cells (Fig. S1 C-D and Movies 3 and 4). IGR1-RAC1^P29S^ melanoma cells had thin, veil-like protrusions all over the cell body, which extended and retracted dynamically, sometimes capturing Jurkat T cell targets (Movie 3), which accumulated within the melanoma cells over time (Fig. S1C). By contrast IGR1-RAC1^WT^ cells were far less dynamic, with lamellipodial activity primarily at the poles of spindle-shaped cells. IGR1-RAC1^WT^ did not accumulate Jurkat T cells during the imaging time (Fig. S1D, and Movie 4). IGR1-RAC1^P29S^ extended membrane protrusions that wrapped around Jurkat T cells (Fig. 1K-K’ and Movie 5), internalizing the target cell within minutes (Movie 5). These data indicate that IGR1-RAC1^P29S^ melanoma cells engulf necrotic targets by extending protrusions and forming a phagocytic cup.

### RAC1^P29S^ enhances the engulfment of whole living T cells and cell fragments

In addition to dead cells, metastatic melanoma cells engulf living lymphocytes ^2^. We next tested if RAC1^P29S^ enhances the engulfment of live cell targets. We co-cultured melanoma cells with live Jurkat T cells and assessed engulfment frequencies in fixed samples after 1 day. Both IGR1-RAC1^P29S^ and IGR1-RAC1^WT^ cells contained some Jurkat T cell material (Fig. 2A-C). However, IGR1-RAC1^P29S^ cells engulfed significantly more whole cells than IGR1-RAC1^WT^ cells (Fig. 2D). We conclude that RAC1^P29S^ enhances phagocytosis of whole, living T cells.

**Figure 2.**
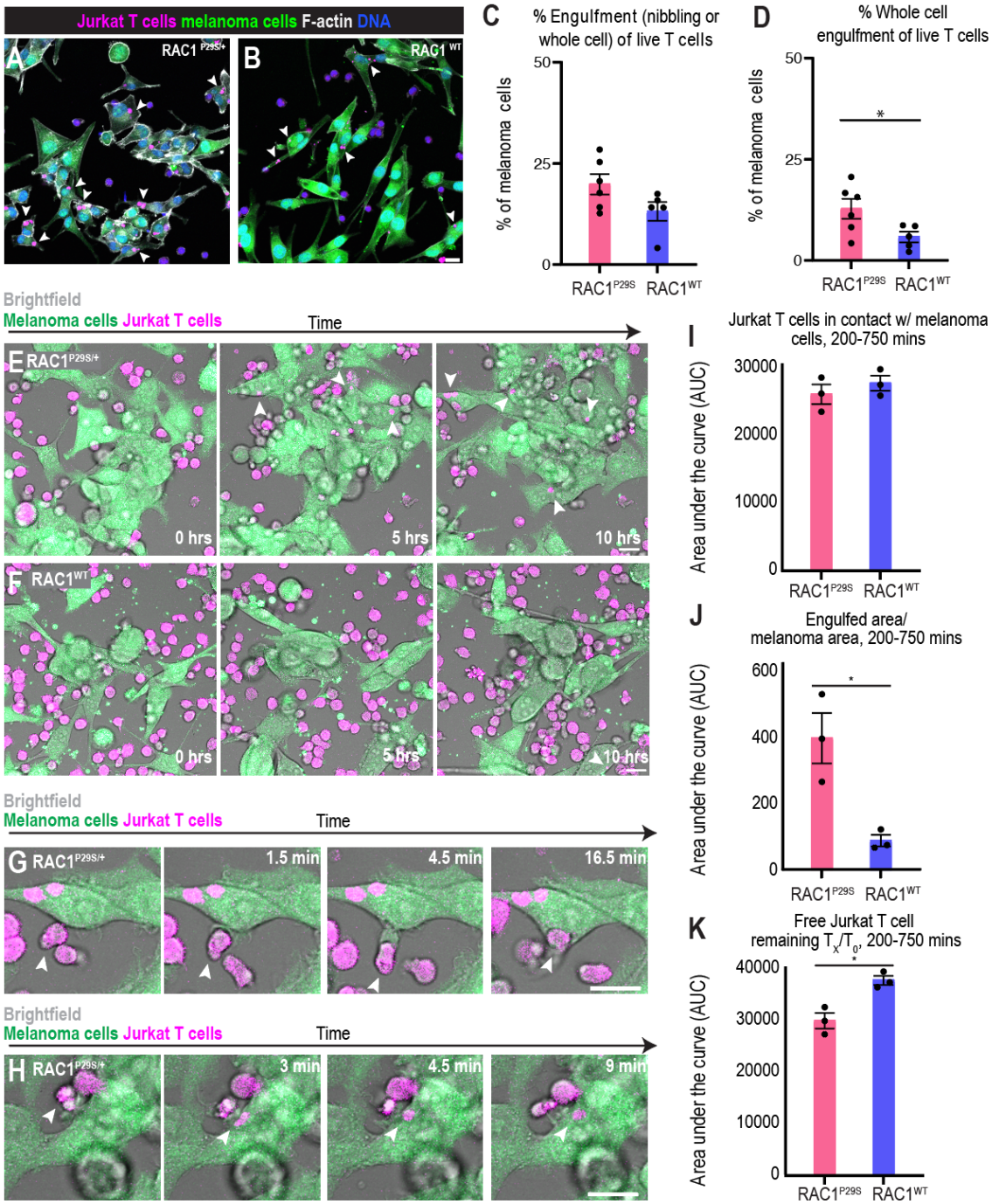
RAC1^P29S^ promotes engulfment of cell debris and pieces of and whole living T cells. A-B Confocal images of melanoma cells labeled with CellTrace-CFSE(green) incubated with 5X living Jurkat T cells labeled with Celltrace-Far Red (magenta). Cells were stained with Hoechst (blue) to label nuclei and Phallodin to label (F-actin). A shows IGR1 cells with the endogenous RAC1^P29S^ mutation. B. shows IGR1 revertant line with RAC1^WT^. Scale bar: 20 microns. C–D. Plots showing the mean (column) +/-standard error of the mean (SEM, black bars) percentages of melanoma cells that engulfed pieces of, or whole, living Jurkat T cells (C) as measured by the presence of Celltrace-Far Red inside the melanoma cell or a whole cell (D) as measured by the presence of Cell-trace overlaid with Hoechst inside the melanoma cell for the indicated conditions. Each dot represents an experimental replicate. N=6 experimental replicates for RAC1^P29S^ and N=5 for RAC1^WT^. A student’s T test was performed. C was not significant. D, *= p<0.05. E-F. Confocal images from a timelapse series of melanoma cells labeled with Celltrace-CFSE (green), cultured with living Jurkat T cells labeled with Celltrace-FarRed. G. Magnified image of RAC1^P29S^ melanoma cell engulfing a full living T cell. H. Magnified image of RAC1^P29S^ melanoma cell engulfing a piece of a Jurkat T cell. Scale bars: 20 microns. I-K. Plots showing the mean (center line) +/-SEM (black bars) of the Area under the curve from 200-750 minutes for measurements of percent of unengulfed Jurkat T cells that are in contact with melanoma cells (I), Engulfed Jurkat area divided by the green cell area (J), or the percentage of remaining Jurkat T cells relative to the initial number of T cells (K). The mean of each experimental replicate is shown (black dots) N=3 experimental replicates. n=10 movies. A Welch’s T test was performed to test for statistical significance *p<0.05.

To gain insight into the dynamics of melanoma cell engulfment of Jurkat T cells over time, we performed live confocal imaging of co-cultures right after adding living Celltrace-FarRed labeled Jurkat T cells and imaged the cells for 12.5 hours. In IGR1-RAC1^P29S^ cells, Jurkat T cells accumulated inside the melanoma cells over time (Fig. 2E, Movie 6). In contrast, and despite extensive interactions between IGR1-RAC1^WT^ cells and living Jurkat T cells, we did not observe many Jurkat T cells accumulate within melanoma cells (Fig. 2F, Movie 7). IGR1-RAC1^P29S^ cells engulfed whole living T cells (Fig. 2G, Movie 8), dead cells (Fig. S2A), and cell fragments attached to Jurkat T cells (Fig. 2H, Movie 9). We also observed living Jurkat T cells, upon prolonged contact with IGR1-RAC1^P29S^ melanoma cells, undergo morphological shrinkage and blebbing typical of apoptotic cells followed by the melanoma cell engulfing T cell fragments (Fig 2H, Fig S2B, Movie 10). Thus, IGR1-RAC1^+/P29S^ melanoma cells appear to kill and then eat T-cells, in addition to consuming live cells.

Living Jurkat T cells have high background pHrodo fluoresce without engulfment, presumably due to their acidic intracellular pH. So, to measure engulfment over time in co-cultures with living T cells, we segmented green-fluorescent melanoma cells and red fluorescent Jurkat T cells from confocal live imaging. We then quantified the objects that were free in the culture, in contact with melanoma cells or fully engulfed by melanoma cells over time (Fig. S3A-E, Movies 11 and 12). The analysis revealed that around 75% of Jurkat T cells were in contact with melanoma cells, and that both genotypes interacted with T cells at similar frequencies (Fig. 2I, Fig. S3C). IGR1-RAC1^P29S^ melanoma cells engulfed more material over 12.5 hours (Fig. 2J, Fig. S3D), consistent with fixed imaging that measured engulfment after 24 hours(Fig. 2A-D). Further, the number of free Jurkat T cells decreased more in IGR1-RAC1^P29S^ co-cultures than in IGR1-RAC1^WT^ co-cultures (Fig. 2K, Fig. S3E). We conclude that RAC1^P29S^ enhances engulfment of living cells, dying cells, and apoptotic debris.

### MERTK levels correlate with cannibalism across a panel of patient-derived RAC1^P29S^ cell lines

We next investigated phagocytosis in additional patient-derived RAC1^P29S^ metastatic melanoma cell lines (Table 1) compared to a control A375 line, which is derived from a primary melanoma tumor and is RAC1^WT^. Less than 10% of A375 cells engulfed heat-killed Jurkat T cells (Fig. 3A) compared to 30-50% in three RAC1^P29S^ lines (Fig. 3B-C). The YUHEF cell line had the highest rates of cannibalism (Fig. 3B-C). To test for engulfment of living targets, we co-cultured the melanoma lines with live Jurkat T cells. Similar trends held: YUHEF and IGR1-RAC1^P29S^ engulfed significantly more living cells than lines expressing RAC1^WT^ (Fig. 3D). One RAC1^P29S^ line, WM1791c, was exceptional in that it exhibited minimal engulfment in either co-culture condition (Fig. 3C-D), so we investigated possible explanations.

**Table 1.** Melanoma Cell lines.

| Cell line | RAC1 mutation status | Tumor stage and origin | Other mutations |
|---|---|---|---|
| A375 | WT | Malignant, skin | BRAF GOF |
| IGR1-RAC1<sup>WT</sup> | WT | Metastasis, lymph node | BRAF GOF |
| IGR1-RAC1<sup>P29S</sup> | P29S | Metastasis, lymph node | BRAF GOF |
| YUHEF | P29S | Metastasis, lung | NF1 LOF MERTK GOF |
| WM1791c | P29S | Metastasis, site unspecified | KRAS GOF |
| WM3060 | P29S | Metastasis, left axillary node | NRAS GOF |
| SKMEL2 | WT | Metastasis, thigh | NRAS GOF |
| SKMEL5 | WT | Metastasis, axillary lymph node | BRAF GOF |
| SKMEL28 | WT | Metastasis, lymph node | BRAF GOF |
| YUROB | WT | Metastasis, abdomen | HRAS GOF |

GOF: Gain of function LOF: loss of function.

**Figure 3.**
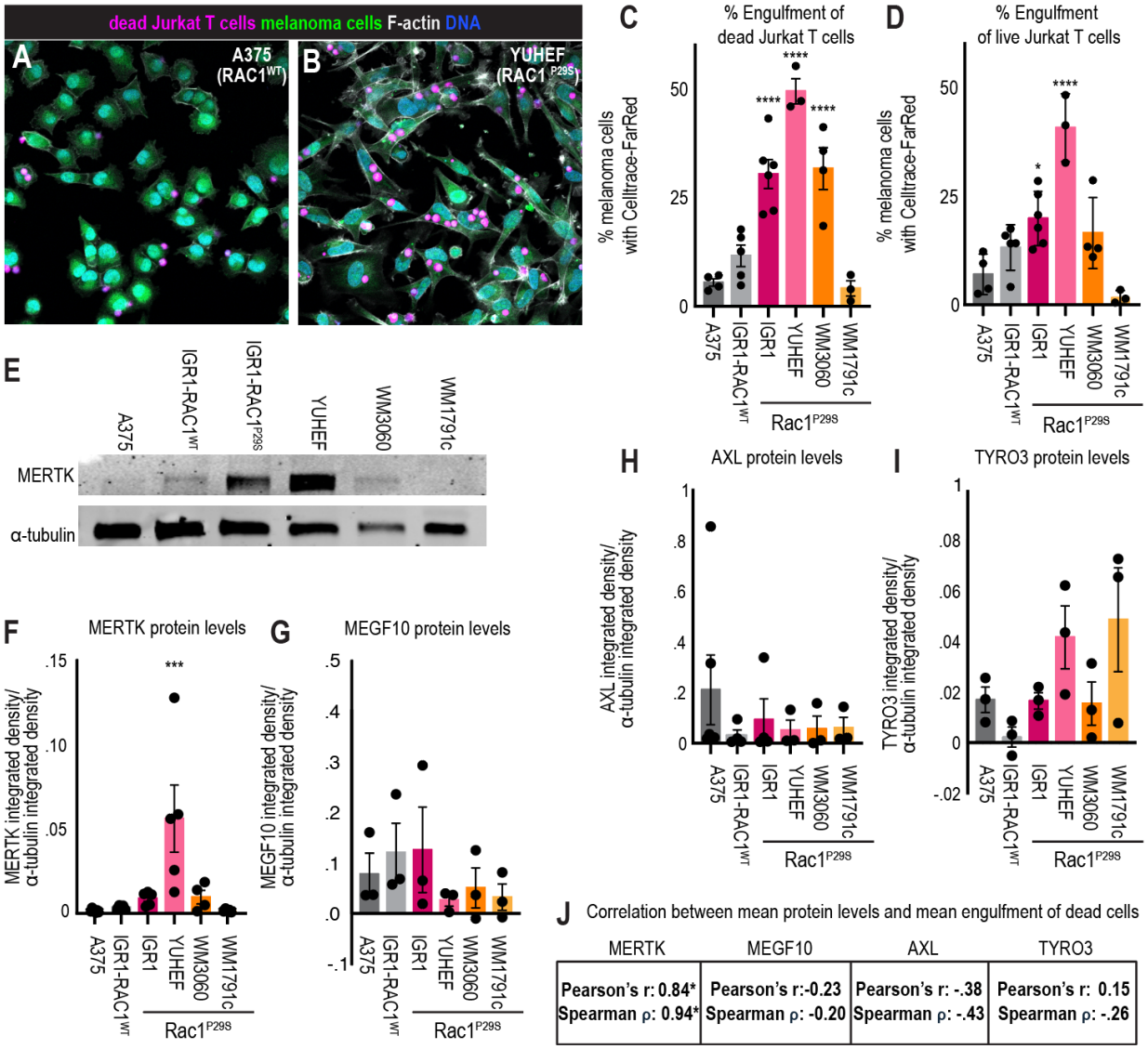
RAC1^P*29S*^ lines display distinct rates of cannibalism that correspond with MERTK levels. A-B. Confocal images of melanoma cells labeled with CellTrace-CFSE(green) incubated with 5X heat-killed Jurkat T cells labeled with Celltrace-Far Red (magenta). Cells were stained with Hoechst (blue) to label nuclei and Phallodin to label (F-actin). A shows A375, the control RAC1^WT^ line while B shows YUHEF, a RAC1^P29S^ melanoma cell line. C-D. Plots showing the mean (column) +/-standard error of the mean (SEM, black bars) percentage of melanoma cells that engulfed a heat-killed Jurkat (C) or live Jurkats (D) as measured by the presence of Celltrace-Far Red inside the melanoma cell for the indicated conditions. Each dot represents an experimental replicate, N≥3 experimental replicates. The percent engulfment was compared to the A375 control by a one way ANOVA followed by a Dunnetts multiple comparison test. *p<0.05. ****p<0.0001. E. Western blots of protein lysates from the indicated cell lines and probed for MERTK as well as the loading control, alpha-tubulin. F-I. Plots showing mean +/-SEM integrated density normalized to the loading control for MERTK (F), MEGF10 (G), AXL (H), and TYRO3(I) in the indicated cell line.s Protein levels were compared to the A375 control by a one way ANOVA followed by a Dunnetts multiple comparison test ****p<0.0001. Each dot represents an experimental replicate (N≥3). J. Pearson’s and Spearman’s correlation values of mean engulfment and mean protein levels of each cell line. *p<0.05

We and others have observed Rac hyperactivation specifically enhances engulfment-receptor-dependent phagocytosis ^16,18^. TAM receptors (<u>T</u>YRO3, <u>A</u>XL, <u>M</u>ERTK) promote engulfment of dead cells by professional phagocytes ^26^. Intriguingly, they are also commonly overexpressed in melanoma ^27^, and MERTK overexpression induces efferocytosis in breast cancer cells ^28^. Notably, WM1791c, the one RAC1^P29S^ line that did not exhibit cannibalistic behavior also lacks detectable MERTK protein (Fig. 3E). We conclude that RAC1^P29S^ alone is not sufficient to induce cannibalism. Rather, RAC1^P29S^ likely enhances receptor-dependent engulfment.

To investigate which engulfment receptors might contribute to melanoma cell cannibalism, we first compared IGR1-RAC1^P29S^ and A375 Depmap RNA expression datasets. We found that the engulfment receptors MERTK and MEGF10 were enriched approximately 7-fold and 10.6-fold, respectively, in IGR1-RAC1^P29S^ compared to A375 (log2 differences of 2.8 and 3.4) (Fig. S4A). The bridging ligand PROS1, which binds phosphatidyl serine and MERTK, was also overexpressed in IGR1-RAC1^P29S^ compared to A375 (Fig. S4A). In contrast, TYRO3 was expressed at similar levels in IGR1-RAC1^P29S^ and A375 whereas AXL was downregulated (Fig. S4A).

To assess protein expression of engulfment receptors across all of the lines, we performed western blotting for MERTK, MEGF10, TYRO3, and AXL on cell lysates and used α-tubulin or GAPDH a loading control (Fig. 3E and Fig. S4B-D). All of the cannibalistic lines had detectable MERTK (Fig. 3E). Quantification of protein levels showed MERTK was higher on average in the three cannibalistic lines with YUHEF, the highest engulfing line, showing the most significant increase (Fig 3F). YUHEF carries a gain of function mutation in MERTK in addition to its overexpression ^29^. Despite elevated mRNA expression, MEGF10 protein was not measurably elevated in IGR1-RAC1^P29S^ relative to A375 or IGR1-RAC1^WT^ and was reduced in the other RAC1^P29S^ lines (Fig. 3G). Consistent with the mRNA analysis, neither AXL nor TYRO3 was consistently elevated in cannibalistic lines (Fig. 3H-I). Of the receptors tested then, only MERTK levels correlated significantly with engulfment frequency (Fig. 3J).

We note that IGR1-RAC1^P29S^ cells express higher levels of MERTK than IGR1-RAC1^WT^ (Fig. 3E and F). This is likely due to RAC-stimulated MERTK transcription because melanocytes independently engineered to express RAC1^P29S^ also exhibit elevated MERTK expression (Fig. S4E). Furthermore, as described below, engulfment in IGR1-RAC1^P29S^ cells was dependent on both MERTK and RAC activity.

### Inhibitors of RAC1, PI3K, branched F-actin polymerization, and MERTK reduce melanoma cell engulfment of dead targets in Rac-mutant cells

To investigate which receptors and signaling pathways are required for cannibalism in RAC1^P29S^ lines, we tested pharmacological inhibitors on two of the RAC1 mutant lines, IGR1-RAC1^P29S^ and YUHEF. We focused on inhibitors of RAC itself and its effectors, transcriptional targets of RAC1 including Myocardin-related transcription factor (MRTF), TAM receptors, and RAS effectors since NRAS and BRAF are commonly activated in melanoma (Fig. 4A). We performed our screen on heat-killed rather than living targets to limit any effect of the inhibitors to the melanoma cell. First, we identified drug concentrations that did not produce significant toxicity compared to DMSO controls over the imaging period (Table 2, methods). Then, we assessed the effects of inhibitors on cannibalism by co-culturing inhibitor-treated melanoma cells with pHrodo-labeled, heat-killed Jurkat T cells over 24 hours (Fig. 4B). IGR1-RAC1^P29S^ cells treated with DMSO showed an accumulation of pHrodo signal at 24 hours (Fig. 4C) whereas cells treated with inhibitors including for RAC1/2/3 (EHT1864), Arp2/3 (CK-666), and MERTK(UNC2025) had reduced levels of pHrodo (Fig. 4D-F). To quantify engulfment, we measured the pHrodo area relative to melanoma cell area and normalized it to the average of DMSO controls for each experimental replicate. Inhibitors targeting MERTK, PI3 kinases, BRAF, MRTF, Arp 2/3, and F-actin polymerization significantly reduced engulfment in both cell lines (Fig. 4G-H). Inhibitors targeting ERK, AKT, RAS and Aurora A reduced engulfment in YUHEF but not significantly in IGR1-RAC1^P29S^, whereas inhibition of AKT increased engulfment only in IGR1-RAC1^P29S^, reflecting cell line differences. Two structurally distinct compounds were tested against several targets (Table 2). Two separate MERTK inhibitors and two F-actin inhibitors reduced engulfment in both lines, whereas for Arp2/3 and MRTF only one of the two compounds reached significance in IGR1 (Fig. 4G), potentially because some inhibitors proved to be toxic and had to be used at lower concentrations (see Methods). Overall, these results support a model whereby dead cell recognition occurs through MERTK while RAC-mediated polymerization of branched F-actin and PI3-Kinase membrane remodeling promote phagocytic cup formation.

**Table 2.**
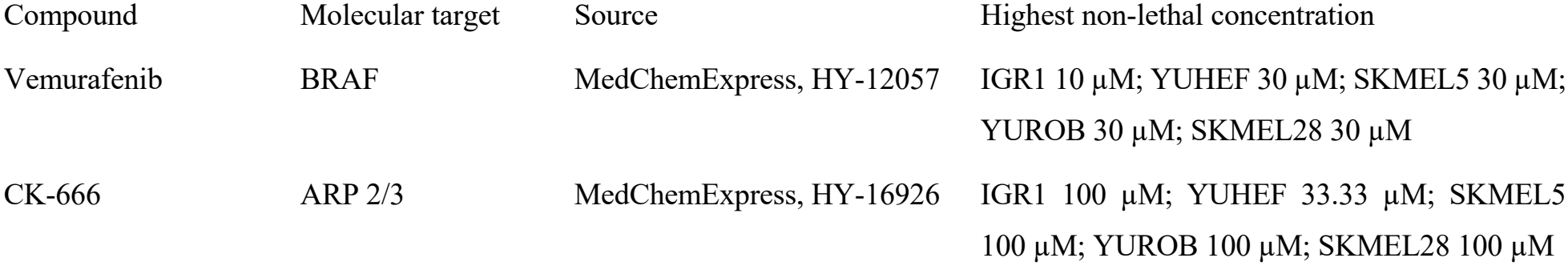
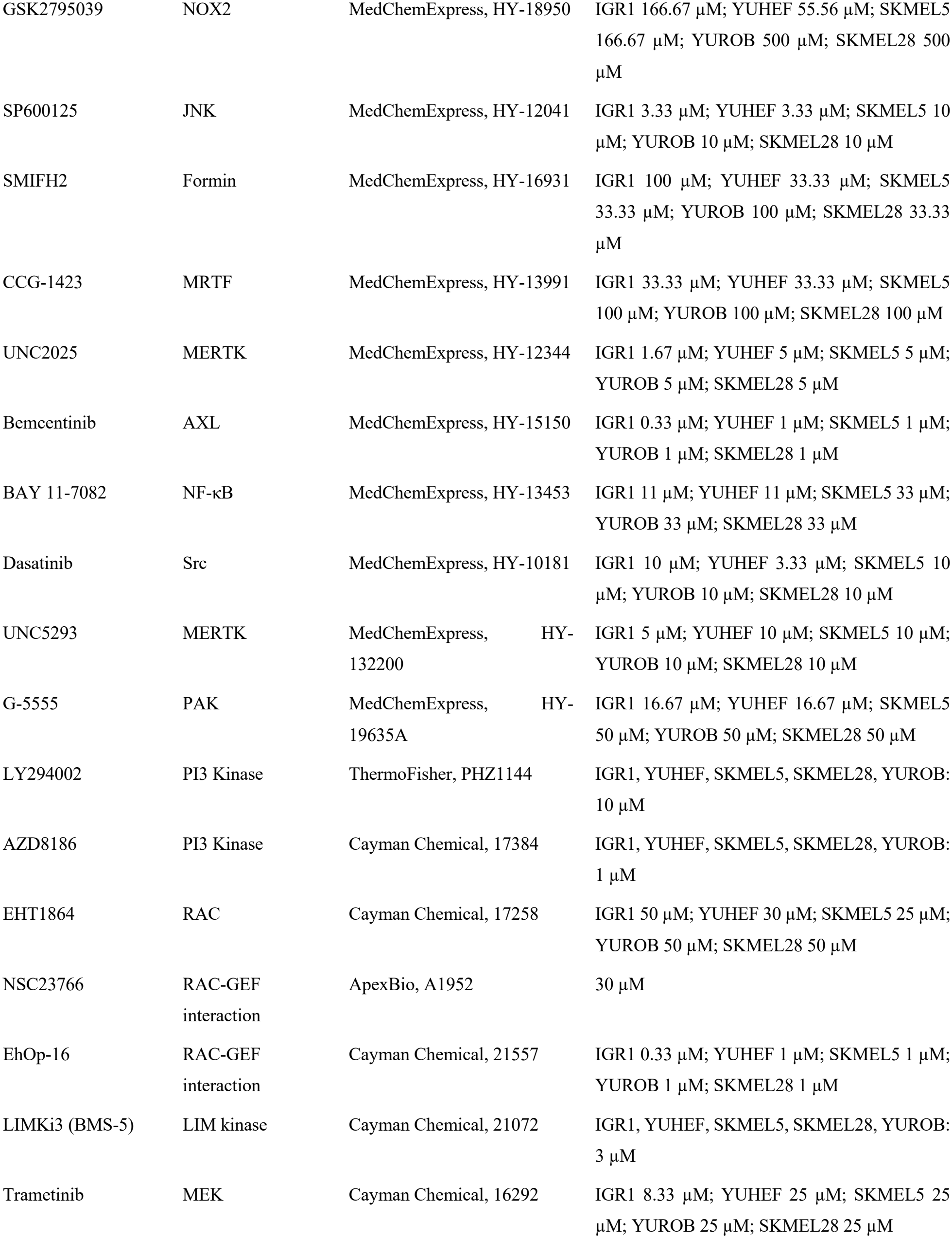
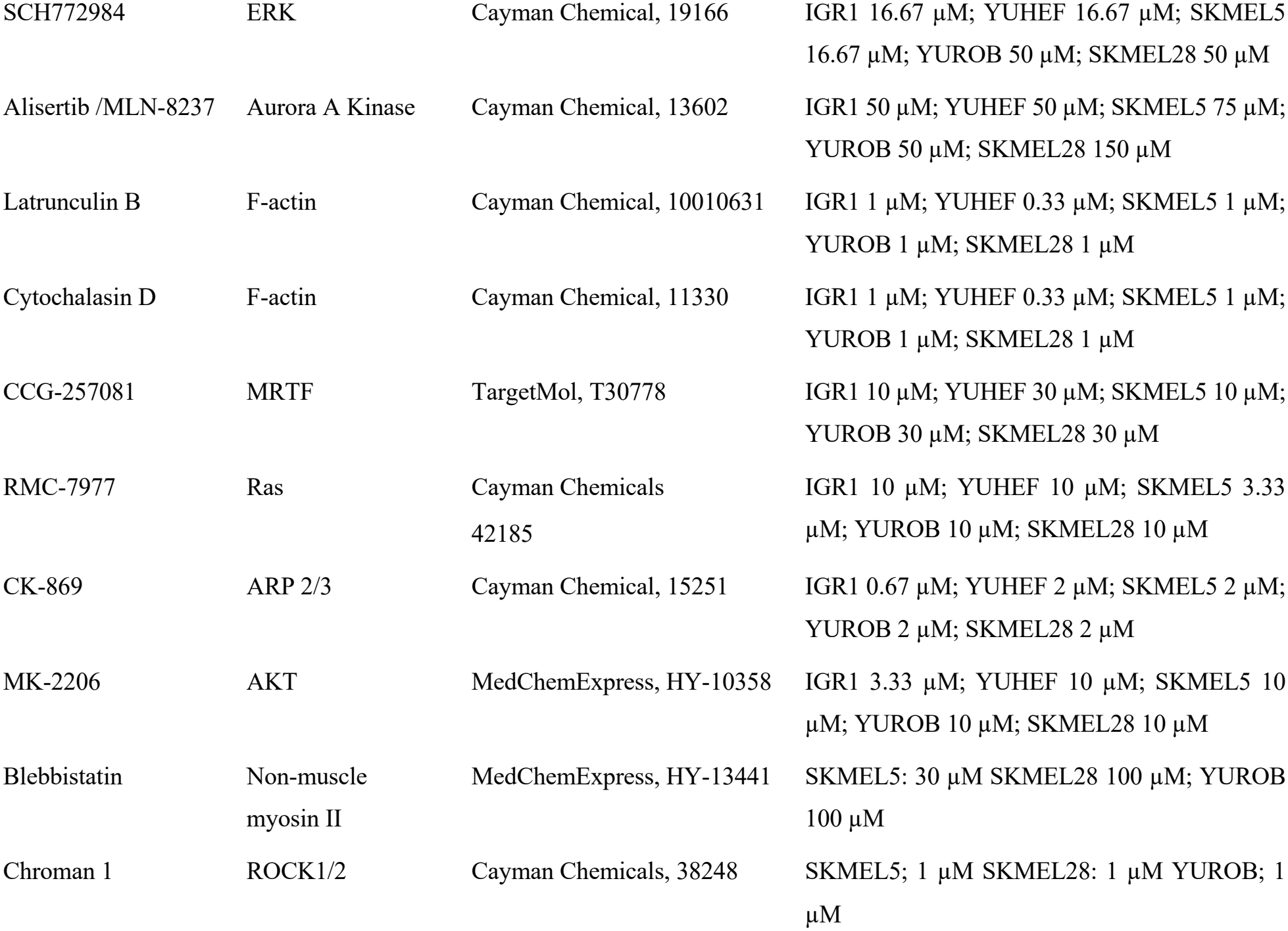
Compounds sources and concentrations.

**Figure 4.**
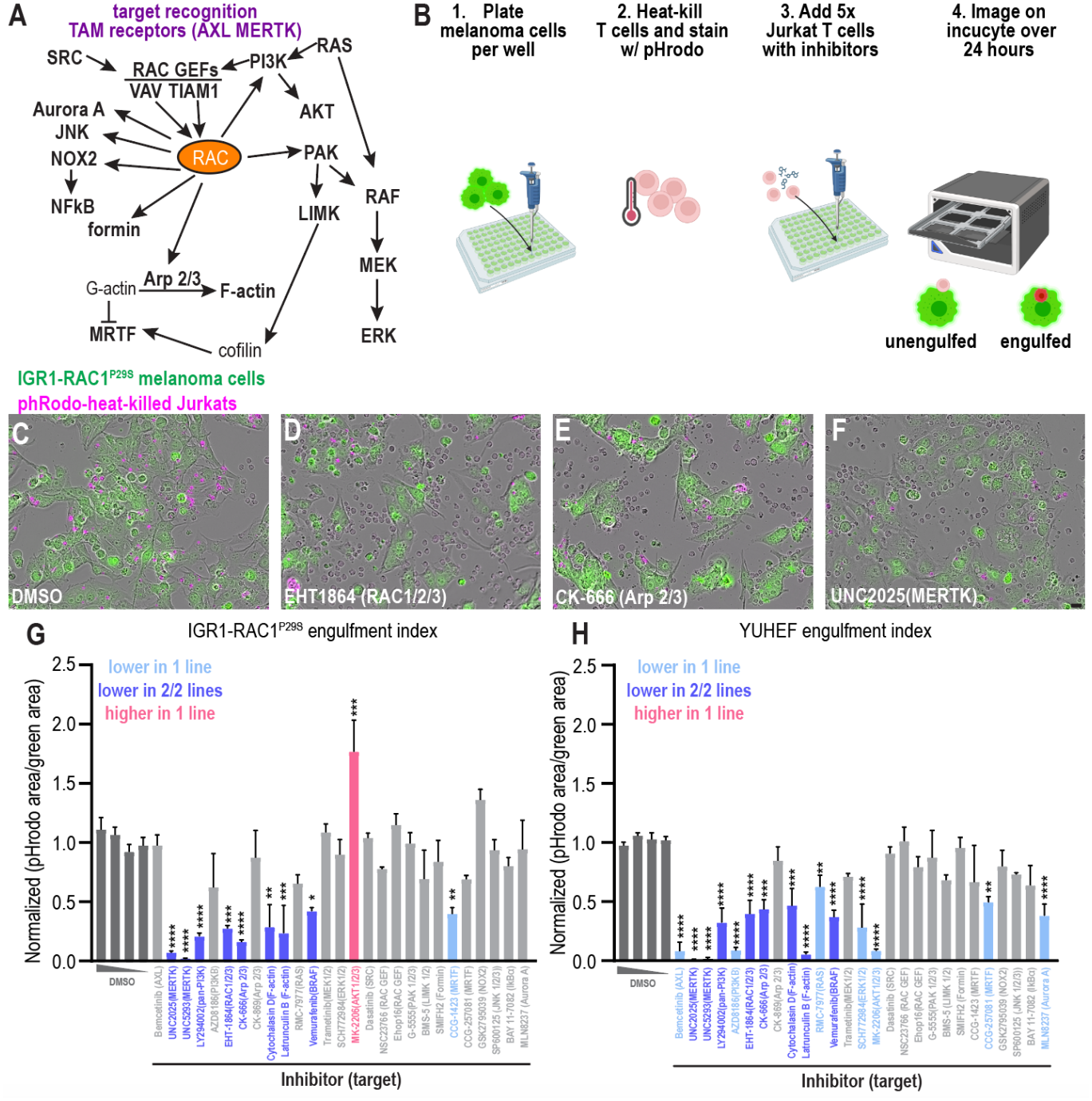
Inhibitors of RAC1, PI3K, branched F-actin, and MERTK reduce melanoma cell engulfment of dead targets in RAC1^P29S^ melanoma cells. A. Schematic of RAC1 and RAS signaling pathways, highlighting the targets for inhibitor screen. B. Schematic showing experimental design of inhibitor screen. C-F. Select fluorescence images of IGR1-RAC1^P29S^ melanoma cells (green, CellTrace-CFSE) co-cultured with pHrodo-labeled, heat-killed Jurkat T cells (magenta), for 24 hours and incubated with the following drugs: DMSO (C), 50 micromolar EHT-1864 (RAC1/2/3 inhibitor) (D), 100 micromolar CK-666 (Arp 2/3 inhibitor) (E), and 1.67 micromolar UNC2025 (MERTK inhibitor)(F). Scale bar: 20 microns. G-H. Plots showing the mean +/-SEM of engulfment index as measured by the mean pHrodo area/green area / pHrodo area/green area of DMSO controls from the same plate. Engulfment index means were compared to the 0.03% DMSO control by a one way ANOVA followed by a Dunnetts multiple comparison test. * p<0.05, **p<0.01, ***p<0.001, ****p<0.0001. Light blue bars: 1/2 lines had a significant reduction in engulfment with the indicated inhibitor. Dark blue bars: 2/2 lines had a significant reduction in engulfment with that inhibitor. Pink bar: a significant increase in engulfment. Gray bars: inhibitors that were not statistically different from the control.

### A subset of RAC1WT melanoma cell lines are cannibalistic and overexpress MERTK

Initial work on melanoma cell cannibalism showed that all metastatic melanoma lines tested were cannibalistic while primary tumor cells were not ^2^. The RAC1^P29S^ mutation only occurs in 5-10% of melanomas ^13^, suggesting that additional mechanisms enhance cannibalism in RAC1^WT^ metastatic melanoma cells. We selected a panel of 4 RAC1^WT^ metastatic melanoma cell lines to test with heat-killed and living Jurkat T cells (Table 1). We compared their rates of cannibalism to A375 as a control. One metastatic line, SKMEL2, engulfed few heat-killed Jurkat T cells (Fig. 5A) while SKMEL5, SKMEL28, and YUROB had significantly higher rates of engulfment compared to A375 (Fig. 5B-C). Similar trends were observed for co-cultures with living Jurkat T cells, however SKMEL28 engulfed about half as many living as compared to heat-killed cells (Fig. 5D).

**Figure 5.**
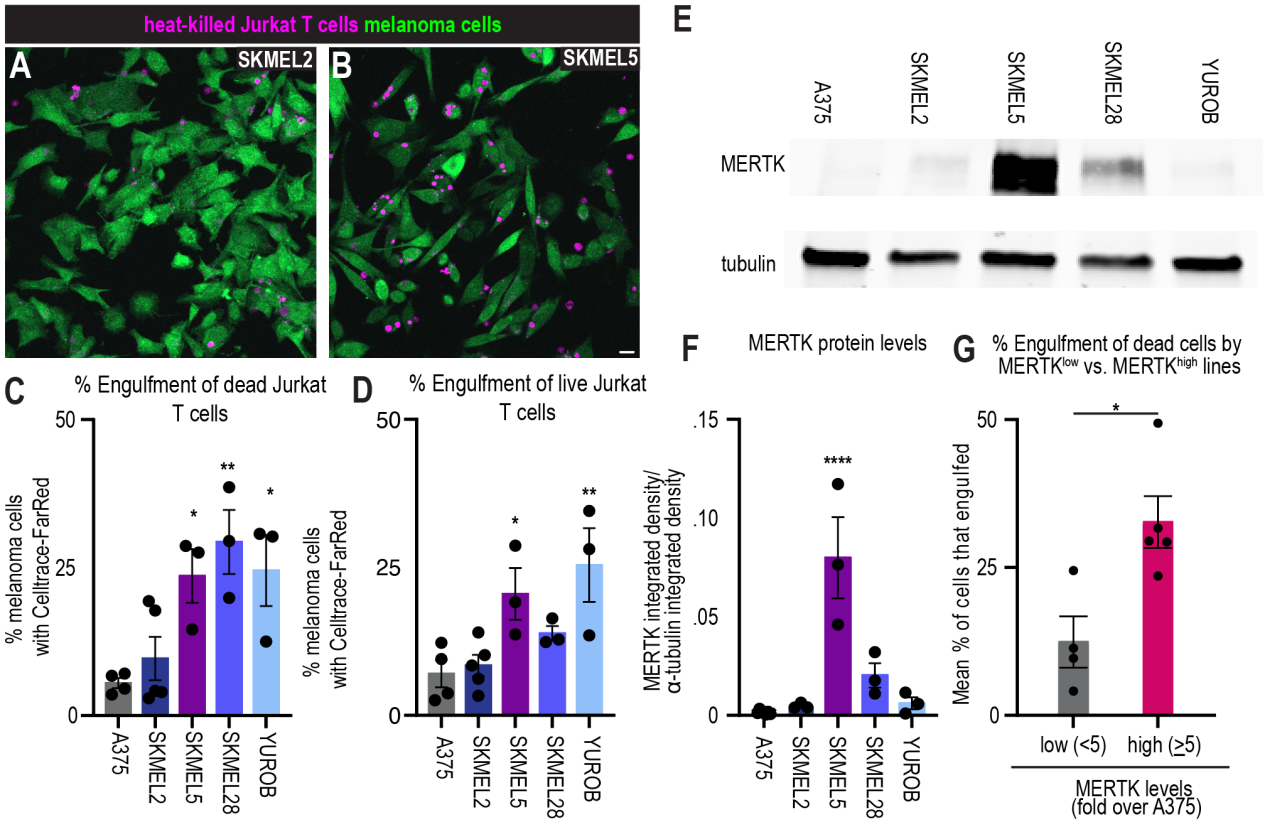
A subset of RAC1^WT^ melanoma cell lines are cannibalistic and overexpress MERTK. A-B Fluorescent images of RAC1^WT^ metastatic melanoma cells stained with CellTrace-CFSE (green) co-cultured with heat-killed Jurkats stained with Celltrace-Far Red (magenta) for SKMEL2 (A), and SKMEL5 (B). C-D. Plot showing the mean (column) +/-standard error of the mean (SEM, black bars) percentage of melanoma cells that engulfed a heat-killed Jurkat (C) or a living Jurkat (D) as measured by the presence of Celltrace-Far Red inside the melanoma cell for the indicated conditions. Each dot represents an experimental replicate N≥3. For C-D, means of the metastatic cell lines were compared to the A375 control by a one way ANOVA followed by a Dunnetts multiple comparison test. *p<0.05, **p<0.01. E. Western blot probed for MERTK and alpha-tubulin for the indicated cell lines. F. Plot showing the mean(column) +/-SEM (black bars) of the integrated density of MERTK divided by the integrated density of tubulin. N≥3. Means of the metastatic cell lines were compared to the A375 control by a one way ANOVA followed by a Dunnetts multiple comparison test.****p<0.0001. G. Plot showing the mean(column) +/-SEM (black bars) of the average engulfment % for each line that was low in MERTK (<5 fold of A375 control) or high in MERTK (≥5 fold of A375 control). An unpaired Student’s T test was performed, *p<0.05.

We next measured MERTK protein expression by western blot and found that SKMEL5 had the highest expression of MERTK followed by SKMEL28 (Fig. 5E-F), consistent with RNA expression data (Fig. S5A). The other cannibalistic line, YUROB, had barely detectable MERTK (Fig. 5E-F). AXL, TYRO3, and MEGF10 exhibited variable expression levels (Fig. S5B-G).

Taking all the melanoma lines analyzed into account, lines with high MERTK (at least 5-fold higher than A375) had significantly higher engulfment than lines with low MERTK (Fig. 5G). MERTK expression and engulfment were significantly monotonically correlated across the 10 lines (Spearman ρ = 0.76, p = 0.01; Pearson r = 0.54, p = 0.11). In contrast, no other receptor or combination of receptors significantly correlated with engulfment (Fig. S5H). The association was evident among both RAC1^WT^ and RAC1^P29S^ lines. MERTK was also necessary for cannibalism: WM1791c carries RAC1P29S yet has no detectable MERTK and the lowest engulfment of any line in the panel, and MERTK inhibition reduced engulfment in every cannibalistic line tested. YUHEF, which combines high MERTK with RAC1^P29S^, engulfed the most targets of any line. Together these observations indicate that MERTK is required for engulfment of dead targets and that RAC1 hyperactivity amplifies this behavior rather than driving it independently.

### Melanoma cannibalism depends broadly on MERTK and PI3K, with cell line–specific contributions of Src, Rac, Arp2/3 and myosin II

To determine if RAC1^WT^ cannibalistic lines use similar or distinct means to engulf target cells compared to RAC1P29S lines, we treated the lines with the highest rates of cannibalism, SKMEL5, SKMEL28, and YUROB, with pharmacological inhibitors. We implemented the pHrodo assay for SKMEL5, but we found that SKMEL28 and YUROB did not have a robust increase in pHrodo intensity after engulfing pHrodo-labeled targets. Instead, we measured engulfment through analysis of confocal images of melanoma cells cultured with CellTrace-FarRed labeled Jurkat T cells. Inhibitors targeting F-actin, MERTK, PI3 kinase beta, Ras and ERK significantly reduced cannibalism by SKMEL28 cells while PAK inhibition significantly increased cannibalism in this line (Fig. 6A). Like SKMEL28, inhibitors that target MERTK, Arp 2/3, F-actin, PI3 kinase, and ERK significantly reduced engulfment in YUROB and SKMEL5 (Fig. 6B-C). A larger subset of inhibitors reduced engulfment in SKMEL5 and YUROB than SKMEL28 including those targeting JNK, formin and other MAP kinase signaling proteins in SKMEL5 (Fig. 6B), and Axl, Aurora A, NOX2 and Src inhibitors reduced engulfment in YUROB (Fig. 6C).

**Figure 6.**
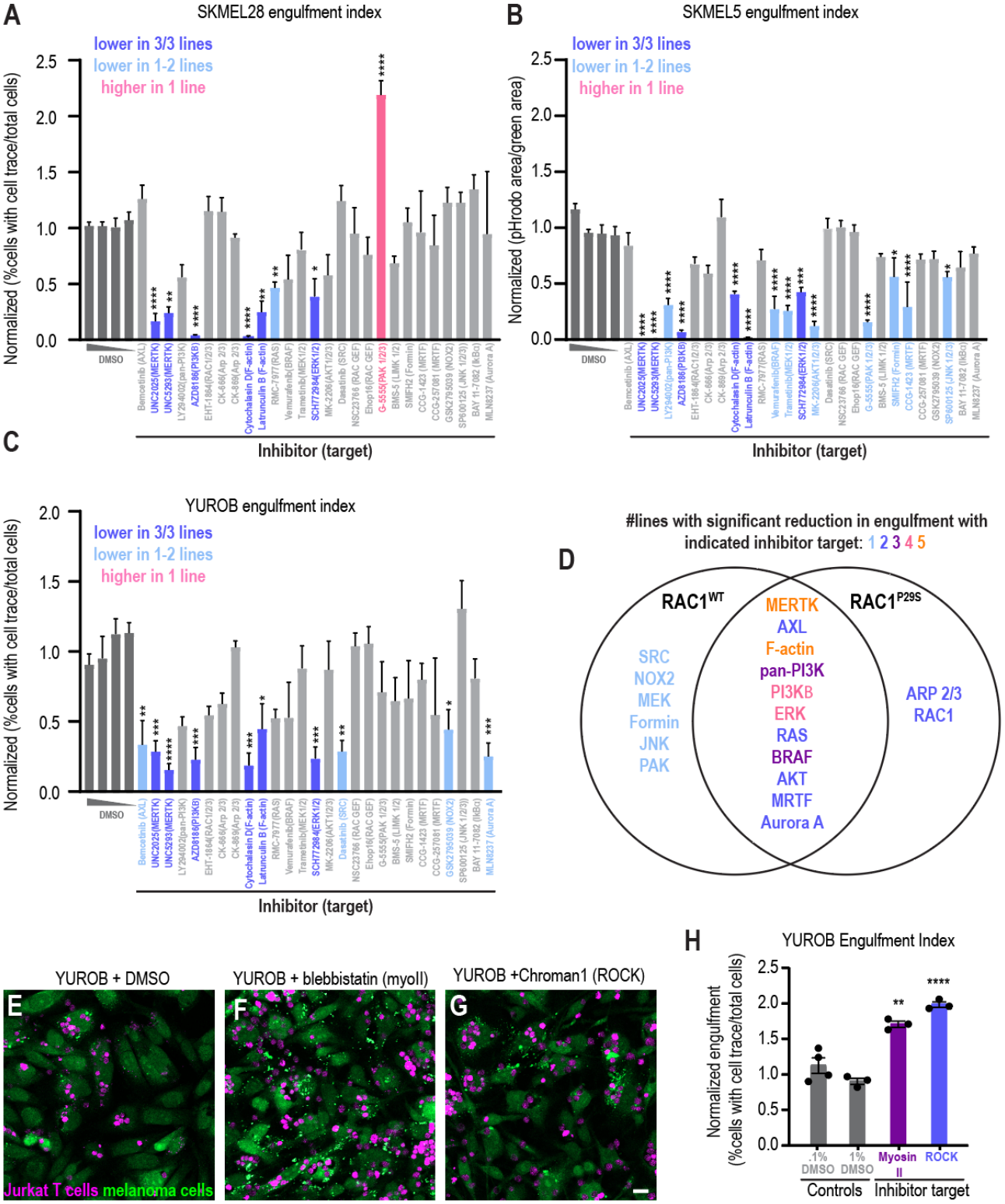
Melanoma cannibalism depends broadly on MERTK, F-actin, and PI3K, with cell line–specific contributions of Src, Rac, Arp2/3 and myosin II. A-C Plot of the means (columns) +/-SEM (black bars) of the engulfment index for (A) SKMEL28, (B) SKMEL5, and (C) YUROB. For A and C, the engulfment index was measured by the percent of cells that engulfed a target/total cells divided by the average percent of cells that engulfed a target/total cells in all DMSO controls from the same plate. For B the engulfment index was measured by the mean pHrodo area/green area / pHrodo area/green area of DMSO controls from the same plate. Engulfment index means were compared to the 0.03% DMSO control by a one way ANOVA followed by a Dunnetts multiple comparison test. * p<0.05, **p<0.01, ***p<0.001, ****p<0.0001. Light blue bars: 1 or 2 of the lines had a significant reduction in engulfment with the indicated inhibitor. Dark blue bars: 3/3 lines had a significant reduction in engulfment with that inhibitor. Pink bar: a significant increase in engulfment. Gray bars: inhibitors that were not statistically different from the control. D. Summary of the targets where an inhibitor had a significant reduction in the engulfment index from the screen. The Venn diagram shows inhibitors that reduce engulfment in RAC1^WT^ (left), RAC1^P29S^(right), or both (middle). The color code indicates how many of the 5 lines had a significant reduction with the target of the inhibitor, orange: 5/5, pink: ⅘, purple: ⅗, blue: ⅖, light blue: ⅕. E-G Confocal images of YUROB cells stained with Celltrace-CFSE(green) co-cultured with heat-killed Jurkat T cells stained with CellTrace-Far Red and incubated with DMSO (E), blebbistatin (F), and Chroman 1 (G). Scale bar: 20 microns. H. Plot showing the mean (column) +/-SEM (black bars) engulfment index for the indicated conditions. The engulfment index was measured by the percent of cells that engulfed a target/total cells divided by the average percent of cells that engulfed a target/total cells in all DMSO controls from the same plate. A One Way ANOVA followed by a Dunnetts test to the .1% DMSO control. **p<0.01, ****p<0.0001.

Across all lines tested inhibition of MERTK and F-actin most consistently reduced engulfment of heat-killed Jurkat cells (Fig. 6A-D). PI-3-kinase inhibition also broadly reduced engulfment (Fig. 6A-D). Inhibition of Arp 2/3 and RAC1 reduced cannibalism in RAC1^P29S^ lines (Fig. 6D). In the RAC1^WT^ lines tested, inhibition of Arp2/3 and RAC1 reduced engulfment in SKMEL5 and YUROB about ∼2-fold but did not reach statistical significance, and had no discernible effect in SKMEL28 (Fig. 6D). Although we cannot rule out the possibility that the need to use different assays contributed to some of the apparent cell line differences, we conclude that cannibalism is broadly sensitive to inhibition of MERTK and F-actin while RAC1 mutant lines are more sensitive to inhibition of RAC-mediated branched actin polymerization.

We next investigated the YUROB line, which is highly cannibalistic despite having both relatively low expression of MERTK (Fig. 5E) and RAC1^WT^. YUROB was uniquely sensitive to Src inhibition (Fig. 6C), and prior literature reports this line has high phosphorylated Src (pSrc) ^30^, consistent with the fact that Src hyperactivity enhances border cell cannibalism of polar cells in Drosophila ^31^. By western blotting, we found high levels of total Src protein in YUROB, however we were unable to confirm elevated levels of phosphorylated Src. Nor did we find any general correlation between Src expression/activity and engulfment across the different lines (Fig. S6A-G). We conclude that while MERTK and RAC appear to be general regulators of melanoma cell cannibalism, additional mechanisms such as Src activation can contribute to this behavior in a context-dependent manner (Fig. 6D).

Compared with RAC1^P29S^ lines, the RAC1^WT^ lines were less dependent on Rac and Arp2/3. We considered two possible explanations. Rho/Rho kinase- and Myosin II-driven contractility drives an alternative and less common “sinking” mode of phagocytosis in professional phagocytes, that is not dependent on Rac-driven protrusive force ^32^ . If melanoma cells employed this mode, myosin inhibition should reduce engulfment. Alternatively, myosin II might oppose engulfment because cortical contractility raises cortical tension, which can be rate-limiting for the uptake of large particles ^33,34^. In that case, myosin inhibition should increase engulfment. To distinguish between these possibilities, we co-cultured the RAC1^WT^ lines with heat-killed Jurkat T cells together with the myosin II inhibitor blebbistatin or the ROCK inhibitor Chroman 1. Relative to the DMSO control (Fig. 6E), both inhibitors increased the frequency with which YUROB cells engulfed dead targets (Fig. 6F-H), while neither significantly affected SKMEL5 or SKMEL28 (Fig. S6H and I). As the targets were dead, the drugs were necessarily exerting their effects on the melanoma cells specifically. We conclude that myosin II is not required for cannibalism in these lines. Rather, in YUROB myosin II restrains it, consistent with cortical tension limiting the engulfment of whole cells in this line, suggesting again that YUROB exploits a distinct Rac-independent mechanism.

Overall, our data show that melanomas have evolved multiple pathways to engulf whole cells but converge on MERTK to engulf dead and living cell targets. Overexpression of MERTK or activation of RAC1 through RAC1^P29S^ mutations are ways by which melanoma cells become phagocytic.

## Discussion

Cell-in-cell structures have been reported in over 25 cancer types since the early 1900s, and are associated with poor prognosis ^3,35,36^. However, the mechanisms controlling their formation are not well understood. Metastatic melanoma cells avidly engulf a wide range of targets, including latex beads, which suggested that cannibalism might proceed without specific recognition of the target cell ^2^. We find instead that melanoma cells require the efferocytosis receptor MERTK to engulf necrotic T-cells.

RAC1 P29 is the third most commonly mutated codon in cutaneous melanoma, after BRAF V600 and NRAS Q61. Here we ascribe a new function to this driver mutation, in addition to its known effects on proliferation, motility, and therapy resistance. RAC1^P29S^ amplifies a latent capacity of melanoma cells to feed on other cells including lymphocytes. So melanoma cells do not only passively evade immune attack but actively consume the attacker. RAC1^P29S^ amplifies cannibalism of both dead and living T cells. Furthermore, live imaging shows melanoma cells extending F-actin-rich protrusions that wrap around a target and internalize it within minutes, a reaching mode of phagocytosis that depends on RAC1 and Arp 2/3 and is insensitive to inhibition of ROCK and myosin II. Our results thus address a longstanding question and show that cannibalism in melanoma relies on a receptor-directed mechanism assembled from the machinery used by professional phagocytes and amplified by a driver mutation not previously associated with phagocytic capacity in this cell type.

### Activating RAC1 mutations enhance engulfment of MERTK-mediated cannibalism

We propose that MERTK provides the “grip” on the target cell, while RAC1-driven branched actin polymerization drives the “gulp”. Necrotic cells expose phosphatidylserine, which MERTK binds through bridging ligands such as GAS6 and PROS1. Ligand-engaged MERTK also initiates PI3 kinase and MAP kinase signaling, and activates RAC1 through VAV1 ^37^, consistent with our observations that both RAC1^WT^ and RAC1^P29S^ lines were sensitive to inhibitors of each of these pathways. RAC1^P29S^ amplifies this output in two ways. It raises MERTK protein levels, evident in the IGR1 isogenic pair, and it independently increases cortical F-actin and the dynamic membrane protrusions that capture targets. RAC1 hyperactivity therefore increases both the number of receptors available to grip a target and the efficiency with which a gripped target is internalized. YUHEF, which combines high MERTK with RAC1^P29S^ engulfs the most targets of any line tested.

Conversely, WM1791c carries RAC1^P29S^ but has no detectable MERTK and was the least cannibalistic line we tested. Hyperactive RAC1 is therefore not sufficient on its own to render melanoma cannibalistic. A receptor/ligand interaction is required to specify the target and RAC hyperactivity enhances the effect. The same dual requirement has emerged in engineered systems where acute RAC activation was not sufficient by itself; rather it enhanced HeLa cell engulfment of dead cells only when combined with the MFG-E8 opsonin domain ^18^. Together these observations point to a general principle for how a cell that is not a professional phagocyte acquires phagocytic capacity.

### Melanoma cells engulf targets by reaching phagocytosis

By live imaging, we found that RAC1^P29S^ melanoma cells extend and retract dynamic, thin, veil-like protrusions around the entire cell body. These projections captured Jurkat T cells and wrapped around them to form a phagocytic cup that internalized the targets within minutes. Reverted RAC1^WT^ cells were far less dynamic and accumulated no targets over the same interval. Earlier electron microscopy of melanoma cannibalism described targets appearing to sink into the melanoma cell rather than being surrounded by an extending cup. Melanoma cells may internalize targets by more than one mechanism with the reaching mode predominating where RAC1 activity is high.

The pharmacology supports this idea. Inhibition of RAC1 or of Arp2/3 reduced cannibalism most strongly in RAC1P29S lines, consistent with branched actin polymerization driving the protrusions that surround a target. Inhibition of ROCK or of myosin II, by contrast, did not reduce engulfment in any of the lines tested, and myosin inhibition increased engulfment by YUROB. In professional phagocytes, Myosin II dependence varies with the phagocytosis mechanism whereby complement receptor-mediated phagocytosis requires Rho, Rho kinase and myosin II whereas Fc gamma receptor-mediated phagocytosis proceeds independently of Rho and Rho kinase ^38^. The insensitivity of melanoma cannibalism to ROCK and myosin II inhibition therefore places it with the RAC1- and Arp2/3-dependent mode of uptake rather than with contractility-driven engulfment. Lowering cortical tension may also reduce the mechanical resistance to extending a protrusion around a target, which would favor engulfment and could account for the increased observed in YUROB following myosin inhibition.

Melanoma cells can apparently achieve phagocytosis by more than one route. YUROB is highly cannibalistic despite expressing little MERTK and carrying a wild type RAC1. YUROB was uniquely sensitive to Src inhibition among the lines tested. Src family kinases function immediately downstream of phagocytic receptors, and Src hyperactivity enhances engulfment in other contexts. These findings suggest that phagocytosis is a latent capacity in the melanocytic lineage, consistent with an earlier report of phagocytosis in melanocytes ^39^, rather than a specialized property restricted to one mutational background. Multiple routes to phagocytic behavior may also contribute to its prevalence in metastatic melanoma.

### Implications for the tumor microenvironment

Consuming other cells may contribute to melanoma survival and spread via multiple mechanisms. For example, melanoma cells deprived of serum die off at significantly lower rates when co-cultured with lymphocytes than when cultured with nondigestable bead targets ^2^, indicating that cannibalism provides nutrients. The tumor core lacks blood supply, becoming hypoxic and nutrient-poor, and many cells die there by necrosis. So, it is plausible that an environment that is both nutrient-limited and rich in the targets MERTK recognizes provides a selective pressure for the ability to engulf necrotic cancer and immune cells to survive.

Along with nutrient scavenging, engulfment of necrotic cells by tumor cells may dampen anti-tumor immune responses. Necrosis ruptures the plasma membrane and releases cell contents including damage associated molecular patterns (DAMPs) that trigger inflammatory cytokines^40^, which recruit innate immune cells to engulf dying cancer cells and acquire cancer antigens. Necrotic cores are nonetheless associated with poor prognosis ^21,40^. Current ideas are that the necrotic core selects for aggressive tumor cells^41^ and a shift of tumor associated macrophages toward immunosuppressive efferocytosis ^42^. Tumor cells that engulf necrotic cells would remove the inflammatory material, converting a potential immune stimulus into a nutrient supply. Phagocytosis of living T cells adds a second, more direct form of local immune suppression.

The same logic extends to the cancer treatment setting. Cancer therapies trigger necrotic death of tumor cells ^43^, and pancreatic tumor cells have been shown to engulf dead cells after chemotherapy and dampen the resulting inflammation^44^. In melanoma, where checkpoint blockade and BRAF inhibitors kill large numbers of cells, cells that escape treatment and are capable of cannibalizing the dead cells would blunt immune responses. In addition to dead cell engulfment, the engulfment of living T cells by melanoma cells may locally suppress immune responses in the tumor microenvironment. Cannibalism may therefore be one route by which a tumor converts therapy-induced cell death into an immunologically silent event.

### MERTK as a molecular target in cancer therapy

MERTK is already an attractive target in melanoma. It is overexpressed in 55% of melanoma cell lines, and it promotes tumor cell survival and growth ^27^, and the inhibitor UNC2025 reduces tumor growth in patient-derived melanoma xenograft models ^45^. In addition to its expression in melanoma cells, MERTK is also expressed on tumor associated macrophages where high expression creates an immunosuppressive environment associated with poor prognosis and resistance to immunotherapy. Reducing macrophage MERTK improved the efficacy of anti–programmed cell death ligand 1 therapy ^46^. Our results add a third mechanism of action for this drug. If melanoma cells use MERTK to consume immune cells, then MERTK inhibition would act not only on tumor cell survival and macrophage polarization but also to prevent tumor cell cannibalism. Each of these mechanisms provides independent benefit and all three argue for pairing MERTK inhibition with immune checkpoint inhibitors.

## Materials and Methods

### Cell lines

Cell lines were kept in a humidified incubator at 37°C with 5% CO2. WM3060 and WM1791c were purchased from Rockland and maintained in Leibovitz L-15 media (Thermo Fisher Scientific, 11415064) with 10% fetal bovine serum (FBS, Genclone, 25-514H). A375, SKMEL2, SKMEL5, and SKMEL28 were purchased from ATCC and cultured in DMEM, high glucose GLUTAMAX (Thermo Fisher Scientific, 10569044) with 10% FBS. YUHEF and YUROB were a gift from Ruth Halaban’s lab (Yale University) and were cultured in Opti-MEM™ I Reduced Serum Medium (ThermoFisher Scientific, 31985070) with 10% FBS. The isogenic pair of IGR1-RAC1^P29S^ and IGR1-RAC1^WT^ were a gift from Gaudenz Danuser lab (UT Southwestern and Institute of Human Biology) and cultured in DMEM, high glucose GLUTAMAX (Thermo Fisher Scientific, 10569044) with 10% FBS. Jurkat T cells were obtained from a lab stock and cultured in RPMI 1640 Medium, GlutaMAX™ Supplement, HEPES (ThermoFisher Scientific, 72400120) with 10% FBS. Cell lines were kept in culture for under 15 passages for experiments. Cell lines were tested regularly for mycoplasma and only lines negative for mycoplasma were included in the study.

### Fixed co-culture and assays

Melanoma cells were stained with CellTrace-CFSE (Fisher Scientific, C34554, 5 μM in 1 mL of 1X PBS) for 15 minutes, washed once in media followed by one wash in 1X Phosphate Buffered Saline (PBS), pH 7.2 (Gibco, 20-012-050). Then, cells were resuspended in media and plated at 10,000 cells per well in a glass bottom 96 well plate(Cellvis, Fisher Scientific, P96-1.5H-N) coated with human fibronectin (10 micrograms/mL in 1X PBS; R&D systems, 1918-FN). The following day, Jurkat T cells were stained with CellTrace-FarRed (Fisher Scientific, C34564, 5 μM in 1 mL of 1X PBS) for 15 minutes, washed once in media followed by a wash in 1X PBS. Cells were then resuspended in the media used for the melanoma cell line and 50 μL containing 50,000 cells was added per well. Melanoma and T cells were incubated together for 1 day and then fixed with 4% paraformaldehyde (Electron Microscopy Sciences, 15711) for 15 minutes RT. Cells were then washed twice in 1X PBS followed by 1X PBS with 0.4% Tween20 (1XPBST) for 15 minutes. Cells were then stained with Hoechst 33342 (1 uM, Invitrogen, H1399) +/-Phalloidin-568 (1:500, Invitrogen A12380) for 30 minutes. Cells were then washed twice in 1X PBS and left in 100 uL of 1XPBS. Plates were imaged on a LSM 800 or LSM 980 confocal microscope at 20x magnification. Four images were taken per well.

### Live imaging assays

Co-cultures were prepared as described for the fixed co-culture assay. Immediately after adding Jurkat T cells, cells were imaged on a Zeiss Laser Scanning Confocal 980 confocal equipped with 5% CO2 and 37 C incubation with a 20x air objective. A single z-slice was acquired every 1.5 minutes over 12.5 hours. 2-4 fields of view were acquired for each experimental replicate.

### Inhibitor screen

Melanoma cells were stained with CellTrace-CFSE (5 μM in 1 mL of 1X PBS) for 15 minutes, washed twice in media followed by 1X PBS and plated at 10,000 cells per well in a tissue-culture treated 96 well plate (Falcon, 353072). The following day, Jurkat T cells were heat-killed at 56°C for 30 minutes to induce primary necrosis ^20^. Jurkat T cells were washed with PBS and then resuspended in 1 mL of 1 micromolar pHrodo in 1x PBS for 30 minutes. 1 mL of 1XPBS with 1% BSA was added directly following incubation. Jurkat T cells were then washed 3 times in PBS to remove excess dye, counted, and 50,000 cells were added to each well at the same time as the inhibitor. Inhibitors were initially added at 100 micromolar, 10 micromolar, and 1 micromolar to identify ranges where toxicity was minimized. Inhibitor concentration was decreased if there was toxicity or increased if the highest concentration had minimal effects. After testing, compounds were added at 3 concentrations with 2, 3-fold dilution of the highest concentration. The highest concentration that did not cause lethality was used as an upper bound for the screen. . Plates were imaged on the Incucyte S3 live-cell analysis system (Green/Red Optical Module, 4647) at 20x magnification. 4 images were taken per well every hour over a time course of 24 hours.

### Cell lysate preparation and Western blot

For each cell line, 300,000 melanoma cells were plated in a single well of a 6-well plate and incubated for 2 days in a humidified incubator at 37°C with 5% CO2. On the day of lysate preparation, media was removed, and cells were then washed with 2 mL of cold 1X PBS. 200 uL of RIPA buffer(ThermoFisher, 89900) supplemented with PhosSTOP™ phosphatase inhibitor cocktail and cOmplete™(Millipore Sigma, 4906845001), Mini, EDTA-free Protease Inhibitor Cocktail (Millipore Sigma, 11836170001) and was added and cells were scraped and collected into a 1.5 mL centrifuge tube. Cells were spun at 12,000 RPM for 12 minutes, and the supernatant was collected in a fresh tube. A sample was collected to measure protein concentration using a Pierce BCA kit (Thermo Fisher Scientific, 23227) and adjusted to the lowest concentration. 6X Laemmli Sample Buffer (Thermo Fisher Scientific, J61337.AD) was added to each sample and lysates were frozen. Prior to running the gel, samples were heated at 95 C for 5 minutes. Precision Plus Protein All Blue Prestained Protein Standards (Bio-Rad, #1610373) was added to the first lane, and an equal amount of the measured protein from the BCA was added to each lane and ran on a 4–20% Mini-PROTEAN TGX Precast Protein Gels (Bio-Rad, 4561096,) Then, samples were transferred to a nitrocellulose membrane(BioRad, 1620115,) for 2 hours at 70 V at 4 C. Membranes were blocked for 1 hour in a blocking buffer (LI-COR Biosciences, 927-60003,) at room temperature, and primary antibodies were added in a 1:1 mixture of 1XTBST: blocking buffer overnight at 4 C.

Primary antibodies used were Mer (D21F11) Rabbit Monoclonal Antibody (Cell Signaling 4319T) 1:1000), MEGF10 Polyclonal Antibody (1:2000, ThermoFisher, PA5-87956), Axl Monoclonal Antibody (7E10) (1:1000, ThermoFisher, MA5-15504),Tyro3 (D38C6) Rabbit Monoclonal Antibody (1:2000, Cell Signaling, 5585), Src (1:1000, Cell Signaling, 2108), Phospho-Src Family (Tyr416) Antibody(1:1000 Cell Signaling, 2101), alpha-Tubulin (DM1A) Mouse Monoclonal Antibody (1:3000 Cell Signaling, 3873S), and GAPDH Polyclonal Antibody (1:5000). The next day, membranes were washed 3X for 10 minutes in 1X TBST and then incubated with secondary antibodies (IRDye 680 Lt DAM, Li-cor 926-68022, and IRDye 800 CW DAR, Li-cor, 926-32213) for one hour. Then samples were washed 3X for 10 minutes in 1X TBST and imaged on a LI-COR imaging system. Images were quantified in FIJI by drawing a region of interests to measure the integrated density of the protein of interest, the loading control (tubulin or GAPDH), and the background of the blot.

### Image Analysis

#### pHrodo engulfment index

To analyze images to measure the engulfment index by the pHrodo area, we used Incucyte analysis software, version 2024B to generate masks for the green(CellTrace-CFSE(melanoma cells) and pHrodo (engulfed Jurkat T cells). Thresholds were set for masks in the software by visualizing the masks for CellTrace-CFSE to capture only melanoma cells and for pHrodo to minimize background fluorescence from heat-killed Jurkats alone while capturing the majority of brighter pHrodo area inside of green cells. The area of pHrodo/green cell area was averaged from 4 images per condition, and the mean of all DMSO concentrations was measured in each plate. Data were normalized to the mean DMSO from the same experimental replicate (same plate).

#### Engulfment in live confocal imaging

To analyze timelapse movies from confocal images, a custom MATLAB script (R2024A, Image Processing Toolbox) was used to segment melanoma cells and T cells and measure the percentage of free T cells, in contact T cells, and engulfed T cells. Two-channel (green, red) time-lapse TIFF stacks were analysed in MATLAB, with the channel layout read from each file’s metadata. For each movie, each channel’s intensities were scaled to 0–1 using that channel’s minimum and maximum across the whole movie, then smoothed with a Gaussian filter (σ = 1 pixel). Uneven red background was removed with a morphological top-hat filter (disk radius 25 µm) before thresholding. Each threshold was calculated once per movie, as the median of the per-frame values, and applied to every frame. Green was segmented as a mask; individual green cells were not separated. The threshold started from Otsu’s method. If the pixels below it still split into two groups separated by at least 12 × the background noise, so that very bright cells had raised the threshold above dimmer cells, the threshold was moved down to that split (repeated up to 5 times). Background level and noise were taken from the darkest peak of the intensity histogram: noise was the robust standard deviation (1.4826 × median absolute deviation) of pixels darker than that peak. The final threshold was background + 0.7 × (threshold ™ background). The mask was then dilated by 1 pixel, closed with a 3-pixel-radius disk, and objects smaller than 100 pixels were removed. Areas expressed per green area used the mask with enclosed holes filled (the green footprint). The red threshold was the background (median intensity) + 6 × background noise (1.4826 × median absolute deviation). Objects had internal holes filled, and objects smaller than 2 µm^2^ were discarded. Touching cells were separated by watershed on the distance transform, splitting only where the neck between two objects was at least 1.5 µm narrower than the objects. Objects still larger than 1.6 × a typical cell area (100 µm^2^) were split again with progressively gentler neck requirements. Red objects that remained oversized or irregular, typically a cell merged with surrounding background, were re-segmented using their own brightness. They were cut at background + 30% of their peak intensity, and the cut was raised in 10% steps (up to 80%) until compact, cell-sized cores remained. Material without such a core was discarded as background. Objects were then kept only if they met all of the following: brightest pixel ≥ 4 × a fixed reference level (4 × background noise above background); solidity ≥ 0.8, for objects ≥ 25 µm^2^, area ≤ 300 µm^2^. Objects of 2–25 µm^2^ were classed as fragments and objects ≥ 25 µm^2^ as cells. To classify red cell states, a free T cell was defined as: no pixel within 2 pixels of the green mask, an in contact (touching) T cell was defined as within 2 pixels of the green mask, but not overlapping it. A T cell was also defined as in contact (but not engulfed) when overlapping the green mask, or with green found around ≥ 30% of the object’s circumference, but not meeting the engulfed criteria. An engulfed T cell was defined as surrounded by green and bright where *Surrounded:* the band 3 pixels wide immediately around the object was divided into 36 sectors of 10° around its centroid, and green had to be present in ≥ 80% of the sectors that the band covered. *Bright:* the object’s peak intensity was ≥ 4 × the same reference level used for the background filter. Surrounded objects that weren’t bright were classed as partial. Segmentation methods were optimized by visual observations of the green and red masks over time with the fluorescent channels.

#### Engulfment in fixed confocal imaging

Engulfment was scored manually for fixed co-cultures. The number of cells that fully internalized a piece of or whole cell labeled with Cell-traceFarRed were counted by assessing if the CelltraceFarRed signal (Jurkat T cells) aligned with the CFSE signal throughout the z-sections to avoid counting cells that were attached or above melanoma cells but not fully internalized by melanoma cells. Then, the number of total cells were counted by counting cells with Hoechst combined with CellTrace-CFSE. For the inhibitor screen, 4 images from each well were scored and for engulfment rates across lines, 4 images from 2 technical replicate wells were scored and combined for each experimental replicate. The number of cells that engulfed divided total cells is reported.

#### Phalloidin intensity measurements

Images were analysed with a custom MATLAB script (R2024A, Image Processing Toolbox). Jurkat targets (far-red) were segmented by thresholding and classed as engulfed or free by their overlap with the IGR1 CellTrace signal. Free Jurkats were excluded from IGR1 cells, and engulfed Jurkat pixels were excluded from phalloidin measurements. Hoechst-stained nuclei were segmented by Otsu thresholding and watershed splitting, and used as seeds. IGR1 cell outlines were obtained from the combined CellTrace and phalloidin signals. Touching cells were separated by a nucleus-seeded watershed along the phalloidin-rich cortex, and outer edges were refined to the phalloidin boundary. Cells smaller than 50 µm^2^ were excluded. Masks were visualized for each image to assess that cell segmentation was optimized. Whole-cell F-actin was the mean phalloidin intensity of each cell; peripheral F-actin was the mean intensity within 1 µm of the cell edge. Both were measured on raw images after subtracting the per-image background (median intensity outside cells). Cell values were averaged per field and fields per experiment for each experimental replicate.

### RNA sequencing analysis

Processed RNA-sequencing data for GSE118343 (Lionarons et al., *Cancer Cell* 36:68–83, 2019) were obtained from the NCBI Gene Expression Omnibus as per-sample RSEM gene-level quantifications (GSE118343_RAW.tar), generated by the original authors using cutadapt 1.9.1, STAR 2.5.2 and RSEM 1.2.31 against the GRCm38 mouse genome with Ensembl annotation release 86. Expected counts for the six libraries — three primary melanocyte cultures from *Tyr-CreER*^+/™^;*Trp53*^F/F^;*Rac1*^LSL-P29S/WT^ mice and two from *Tyr-CreER*^+/™^;*Trp53*^F/F^;*Rac1*^WT/WT^ mice, together with the spontaneously immortalized melan-a line used by the authors as a third wild-type replicate — were assembled into a gene-by-sample matrix. Genes with fewer than 10 reads summed across all six libraries were excluded, leaving 18,388 genes. Differential expression between genotypes was assessed using pydeseq2 0.5.4, a Python implementation of DESeq2, with the design ∼genotype, median-of-ratios size-factor normalization, empirical Bayes dispersion shrinkage, Cook’s distance outlier refitting, Wald tests and Benjamini–Hochberg correction; genes with an adjusted *p* value below 0.05 were considered differentially expressed. Log2 fold changes are maximum-likelihood estimates and were not shrunk. To verify that this pipeline reproduces the original analysis, adjusted *p* values for the four SRF/MRTF target genes in Lionarons Figure 3G were compared with the published values: *Srf* (published 0.0069, here 0.0068), *Actg1* (0.0007, 0.00047), *Cyr61*/*Ccn1* (0.0211, 0.0198) and *Cnn2* (1.54e-18, 4.6e-19). Normalized expression is reported as counts per million or, where indicated, as DESeq2 median-of-ratios normalized counts scaled to the wild-type group mean.

### Data quantification and statistical reporting

Samples were excluded if cells were inviable or if there was a plating error and is noted in primary data files for each assay. The experimental replicates are reported in the figures and figure legends. The middle bars and sample sizes, and statistical tests used are reported in the figure legends. To assess normality, the residuals were plotted on a QQ plot to assess normality visually (inhibitor screen) or a Shapiro Wilk test was performed. Representative images comparing two conditions were acquired and processed in the same manner.

### Data and code availability

The custom MATLAB scripts used for the live-imaging engulfment analysis and the phalloidin intensity measurements are available at GitHub (melanomacellsegmentationandengulfment).

### AI assistance

The custom MATLAB scripts (R2024A, Image Processing Toolbox) described above were written with assistance from Claude (Anthropic), which was also used to format the expression summary tables in Figures S4 and S5, to organize datasheets of data summarized in Figures 1, 2, 4, 6, S1, and S2, and to process RNA sequencing data presented in Figure S4E.

## End Matter

## Acknowledgments

We would like to thank Gaudenz Danuser, Ruth Halaban, and Jonathan Chernoff for melanoma cell lines and helpful discussions as well as Emory Campbell for his technical assistance on this project. The work was supported by the American Cancer Society PF-22-091-01-MM (L.P.) and the U.S. National Institutes of Health 1DP1CA300850 award to DJM.

## Author Contributions

L.P. and D.J.M. conceived of the project. L.P. carried out all of the experiments. M.N. offered advice on the design, execution, and analysis of the drug inhibitor screen. L.P. prepared all of the figures and the first draft of the manuscript. All authors edited.

## Declaration of generative AI

Generative AI was used to develop data analysis code, to interpret data, and to proof read the manuscript.

---

**Figure S1.**
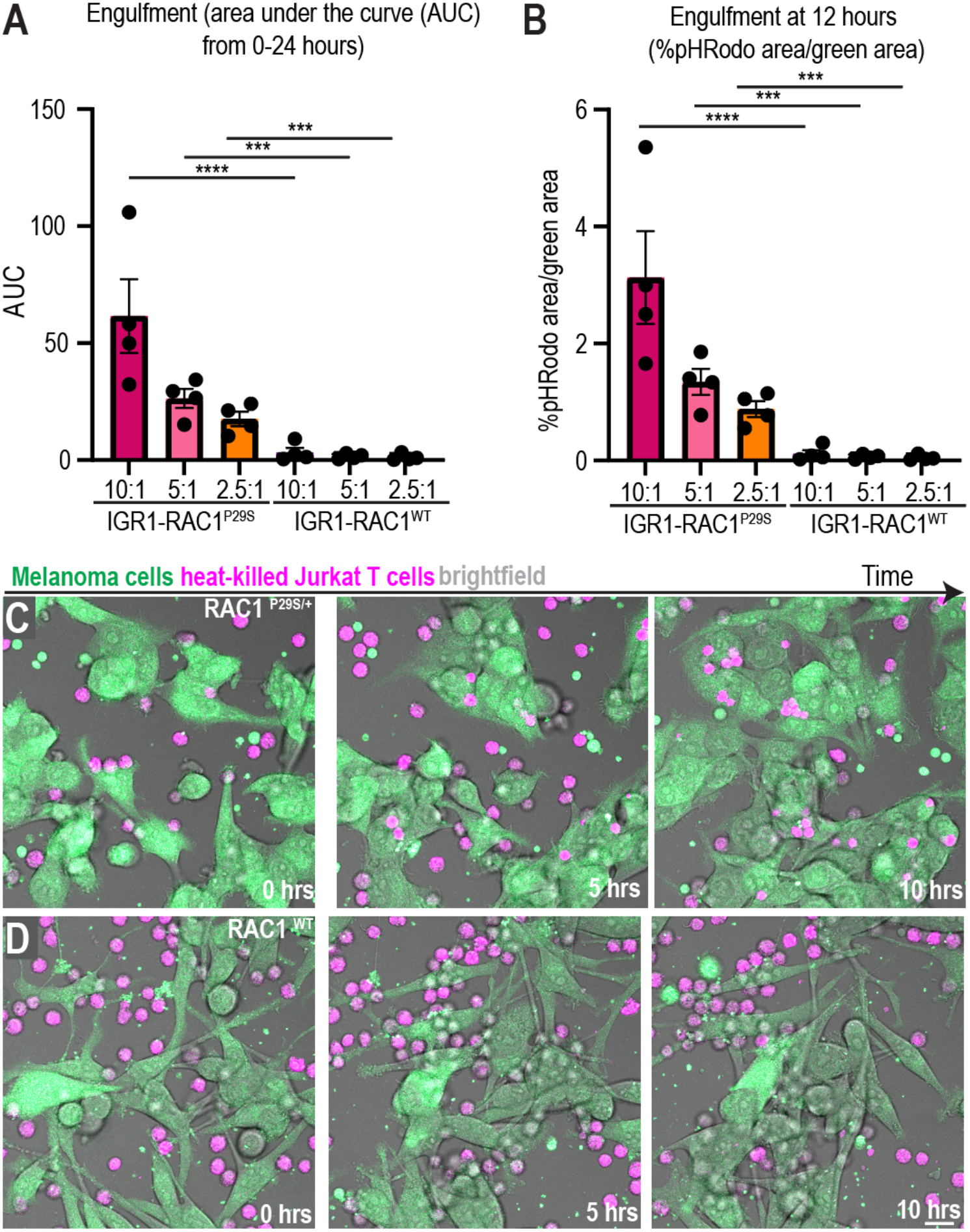
RAC1^P29S^ melanoma cells engulf heat-killed Jurkat T cells within hours. A-B Plots showing the mean +/-SEM area under the curve for 0-24 hours(A) and the mean engulfment at 12 hours (B) as measured by pHrodo area/melanoma cell area for the indicated co-culture conditions . Each dot represents an experimental replicate, N=4. A two way ANOVA was run followed by a Šidák post-tests to determine effects of genotype versus target:effector ratio. ***p<0.001, ****p<0.0001 B. Confocal images from a timelapse series of C. IGR1-RAC1^P29S^ and D. IGR1-RAC1^WT^ cells with melanoma cells labeled with CellTrace-CFSE(green) and heat-killed Jurkat T cells labeled with CellTrace-FarRed (magenta). Time is relative to the start of co-culture. Scale bar: 20 microns.

**Figure S2.**
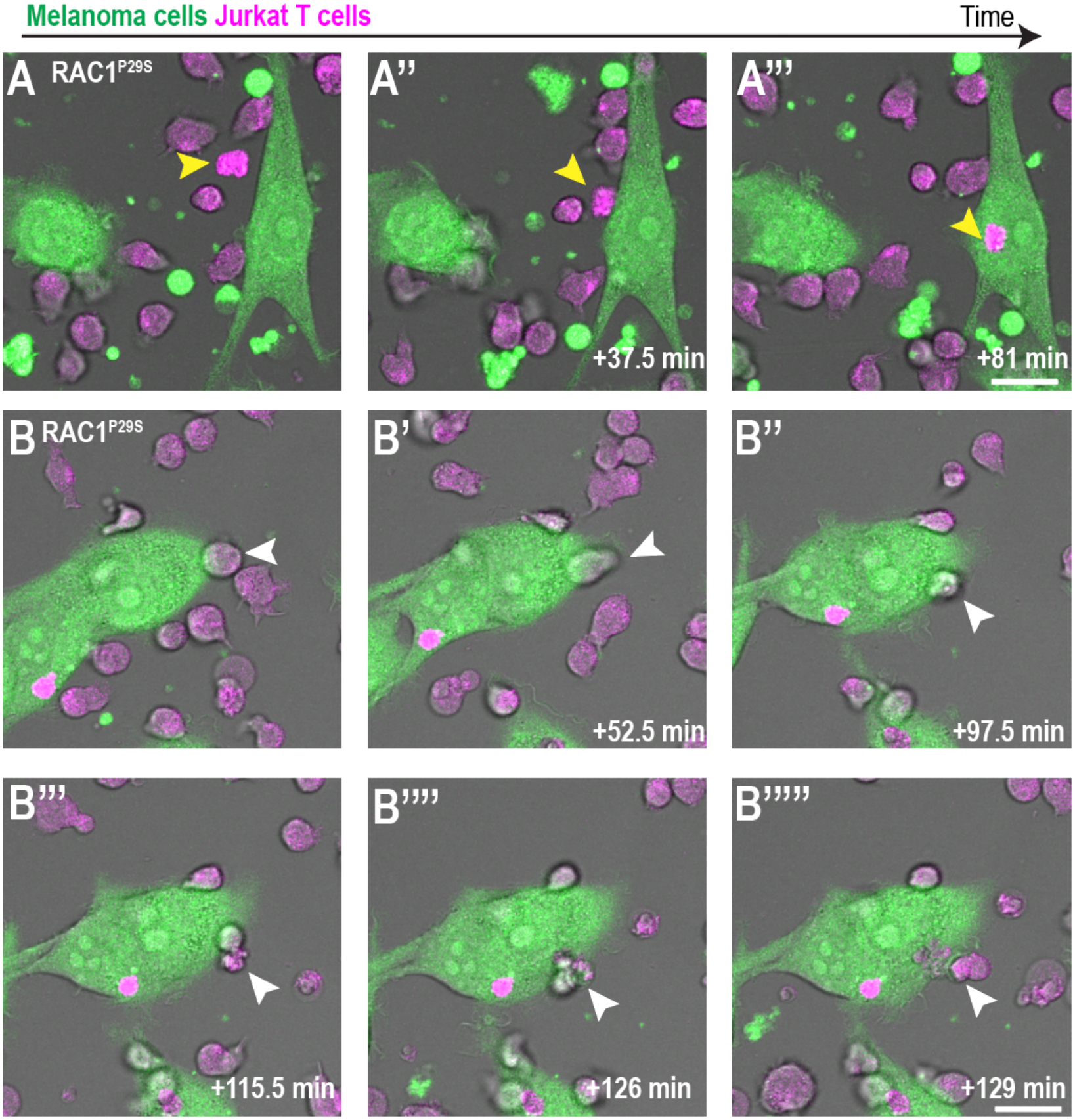
RAC1^P29S^ Melanoma cells engulf dead cells and pieces of cells as they die. **A**. Confocal images showing an example of a IGR1-RAC1^P29S^ melanoma cell labeled with CellTrace-CFSE engulfing a dead Jurkat T cell labeled with CellTrace-FarRed (yellow arrowhead). B. Confocal images showing an example of a IGR1-RAC1^P29S^ melanoma cell labeled with CellTrace-CFSE interacting with a living Jurkat T cell (white arrowhead) that undergoes death and then engulfment by that melanoma cell. Scale bar: 20 microns.

**Figure S3.**
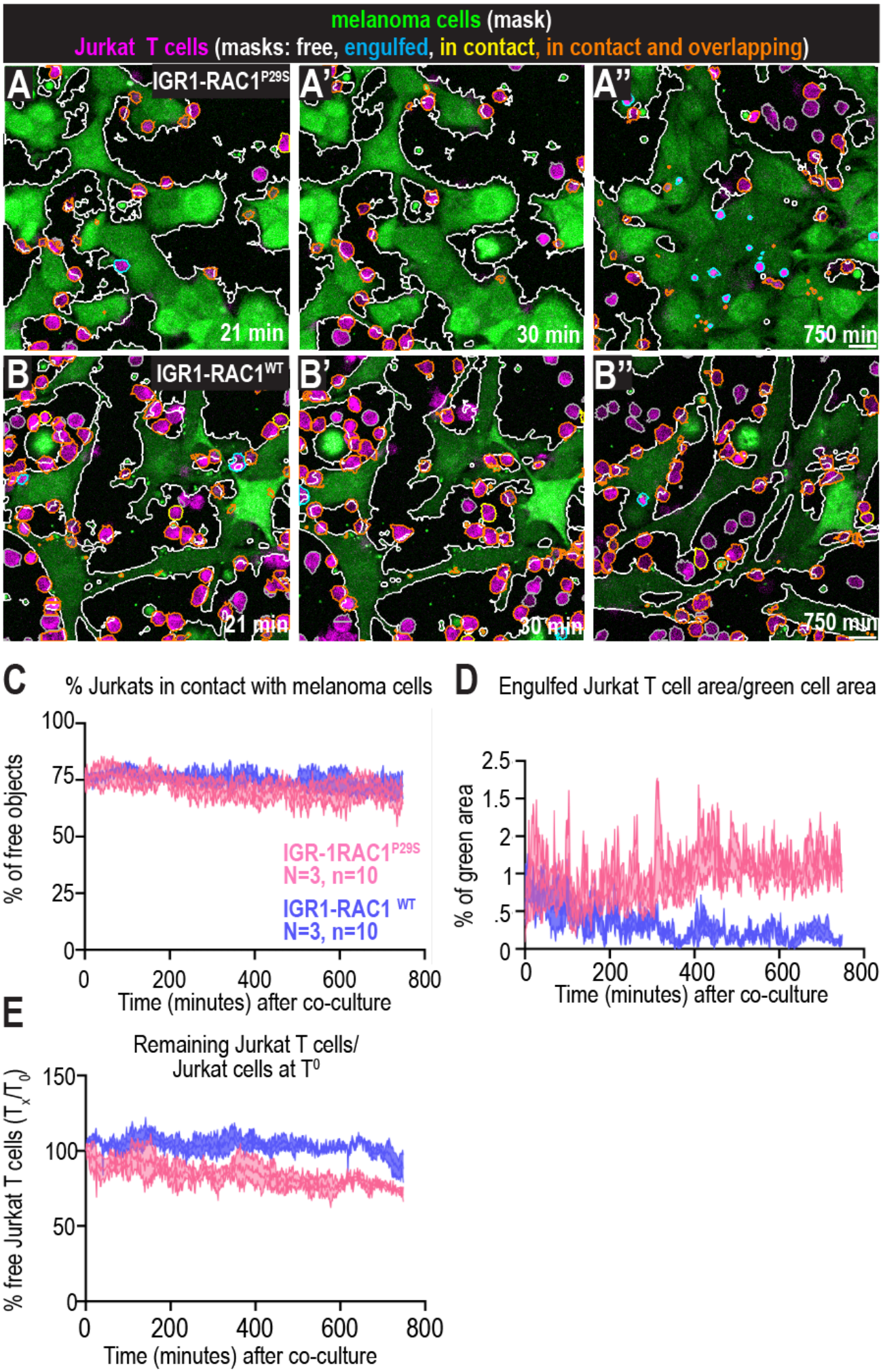
IGR1-RAC1^P29S^ melanoma cells engulf more material than IGR1-RAC1^WT^ in co-cultures with living Jurkat T cells. A-B. Confocal images of Celltrace-CFSE labeled melanoma cells(green) and Celltrace-FarRed labeled Jurkat T cells with overlaying cell masks used for quantification in Figure 2I-K. White outlines mark both melanoma cells and Jurkat T cells. White outlines around Jurkats also mark free cells while yellow and orange outline Jurkat T cells in contact with melanoma cells. Orange indicates partial overlap but not counted as engulfment. Cyan outlines mark cells marked for engulfment. Scale bar: 20 microns. C-E. Plot of mean (center line) +/-SEM (outer lines) of the percent of unengulfed Jurkat T cells that are in contact with melanoma cells over time.

**Fig. S4.**
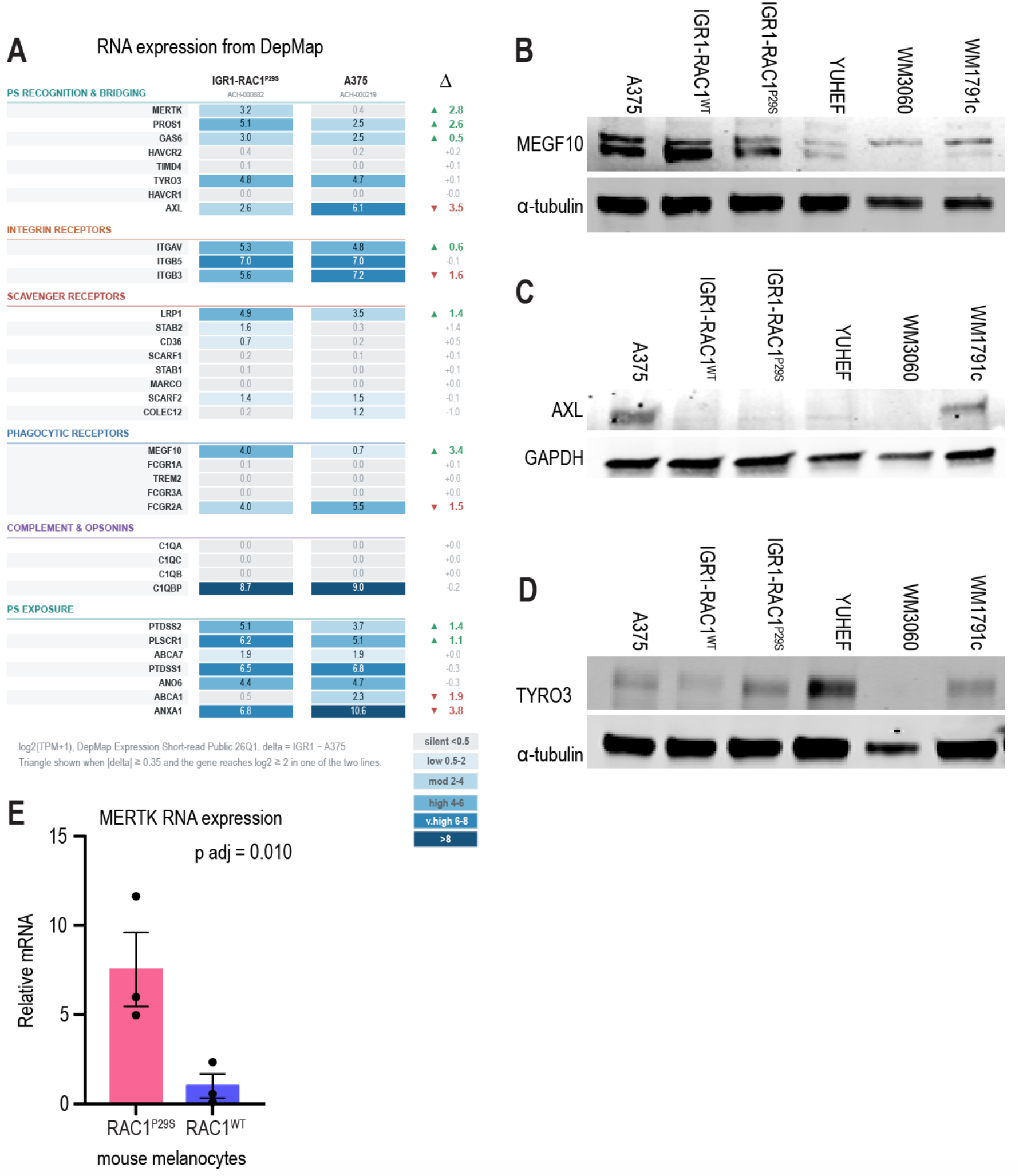
IGR1-RAC1^P29S^ has higher RNA expression of MERTK, PROS1, and MEGF10 than A375. A. Table showing expression data for phagocytosis receptors and associated proteins from Depmap Expression data (Short-read Public 26Q1) for A375 and IGR1 (RAC1^P29S^). The log2(Transcripts per million+1) value is shown. The delta shows the difference between IGR1 and A375 (log2(Transcripts per million+1). The heat map shows the level of expression. B-D Western blots of protein lysates from the indicated cell lines and probed for , MEGF10(B), AXL(C), and TYRO3 (D) as well as loading controls, alpha-tubulin (B,D) and GAPDH (C). E. Plot of mean +/-SEM of relative mRNA expression of mouse melanocytes with the P29S or WT RAC1 gene. Dataset was previously acquired by Lionarons et al., 2019 (GEO accession number: GSE118343). The Benjamini–Hochberg adjusted p value from a DESeq2 Wald test across 18,388 genes is reported.

**Figure S5.**
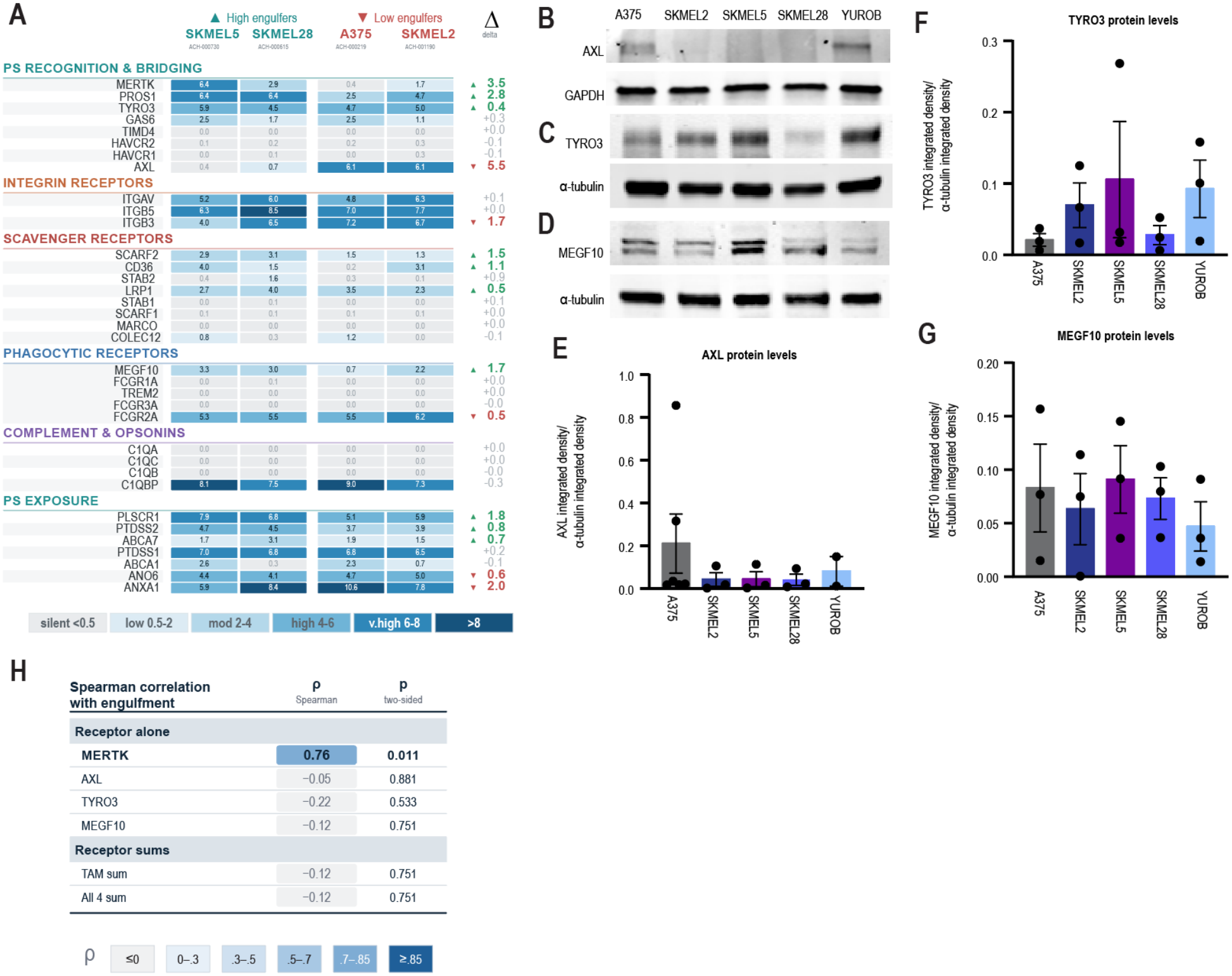
Cannibalistic RAC1^WT^ lines express higher levels of MERTK, and have varying protein levels of other TAM receptors and MEGF10. A. Tables showing expression data for phagocytosis receptors and associated proteins from Depmap Expression data (Short-read Public 26Q1) for high engulfing (SKMEL5 and SKMEL28) and low engulfing (SKMEL2 and A375 lines). The delta shows the difference between high engulfing and low engulfing line mean(SKMEL5+SKMEL28) - mean(A375+SKMEL2). The heat map shows the level of expression. B-D. Western blots of protein lysates from the indicated cell lines and probed for AXL(B), TYRO3(C) and MEGF10 (D) as well as loading controls, GAPDH (B) and alpha-tubulin (C-D). E-G Plots showing mean +/-SEM integrated density normalized to the loading control for AXL (E), TYRO3 (F), and MEGF10 (G) in the indicated cell lines. Each dot represents an experimental replicate (N≥3). Protein levels were compared to the A375 control by a one way ANOVA followed by a Dunnetts multiple comparison test or by a Kruskal-Wallis Test. H. Plot showing spearman correlation between receptor expression and mean rates of engulfment for RAC1^WT^ and RAC1^P29S^ lines.

**Figure S6.**
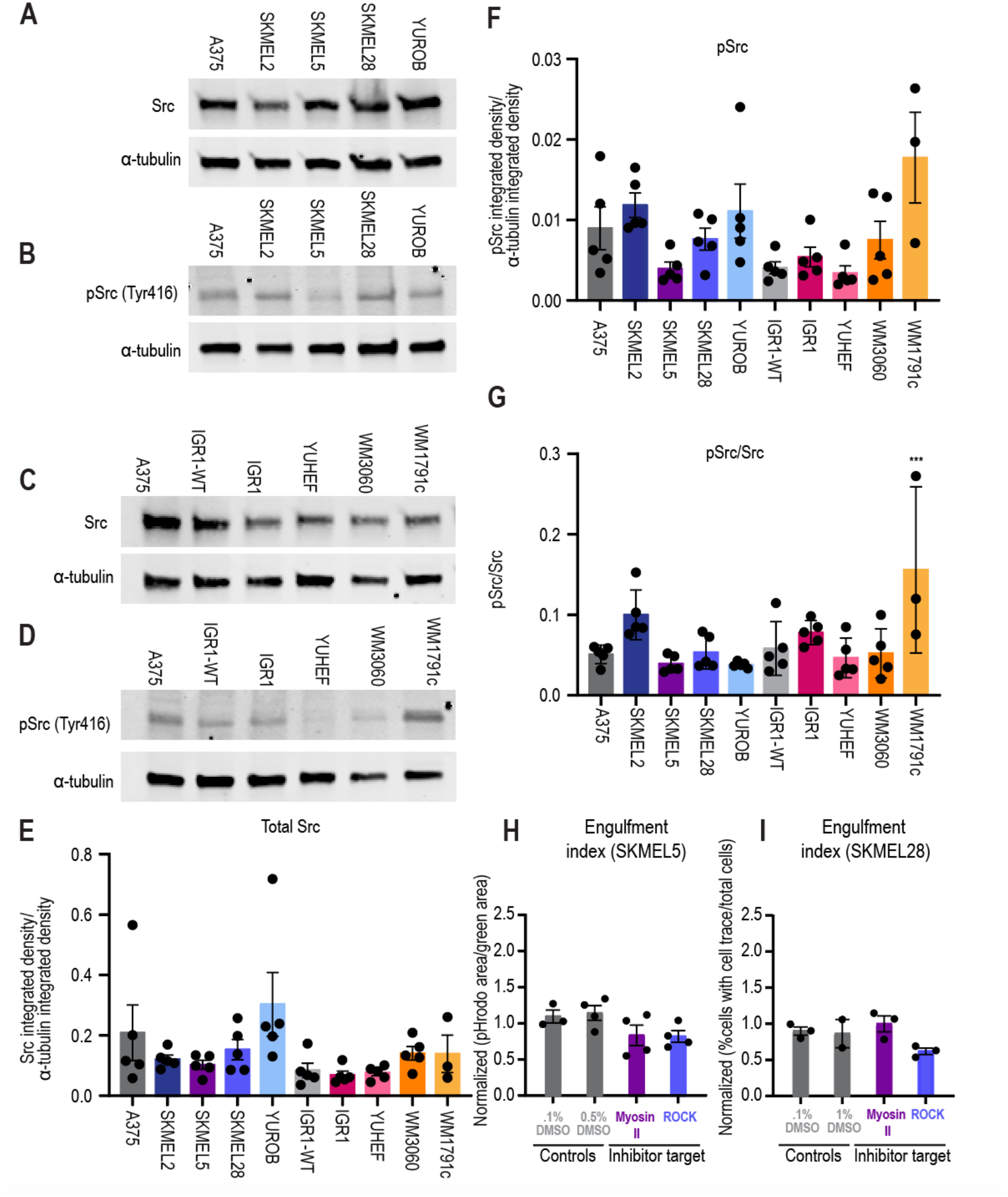
SRC protein expression is high in YUROB while active Src is highest in low-engulfing lines. A-B. Western blots of protein lysates from RAC1^WT^ lines probed for Src (A) and phosphorylated Src at tyrosine residue 416 pSrc(Tyr416) (B) as well as alpha-tubulin as a loading control. C-D. Western blots of protein lysates from RAC1^P29S^ lines probed for Src (C) and phosphorylated Src at tyrosine residue 416 pSrc(pTyr416) (D) as well as alpha-tubulin as a loading control. E-G Plots showing the mean(columns) +/-SEM (black bars) of the integrated density of Src (E), pSrc (F), and pSrc/Src (G) integrated density of the loading control, alpha tubulin. Black dots represent values from individual experimental replicates. Values were compared to the A375 control by One way ANOVAs followed by multiple comparisons Dunnetts tests. ***p<0.001. H-I. Plots showing the mean (column) +/-SEM (black bars) engulfment index for the SKMEL5(H) and SKMEL28(I). For H the engulfment index was measured by the mean pHrodo area/green area / pHrodo area/green area of DMSO controls from the same plate. For I, the engulfment index was measured by the percent of cells that engulfed a target/total cells divided by the average percent of cells that engulfed a target/total cells in all DMSO controls from the same plate. A One Way ANOVA followed by a Dunnetts test to the .1% DMSO control.

## Supplemental Movies

**Movie 1**. IGR1-RAC1^P29S^ melanoma cells (brightfield) co-cultured with pHrodo-labeled, heat-killed Jurkat T cells. pHRodo fluoresces (magenta) when engulfed over time. Time is relative to the start of the co-culture, and the time interval is 1 hour. Playback rate: 5 frames per second (fps).

**Movie 2**. IGR1-RAC1^WT^ melanoma cells (brightfield) co-cultured with pHrodo-labeled, heat-killed Jurkat T cells. pHrodo fluoresces (magenta) when engulfed over time. Time is relative to the start of the co-culture, and the time interval is 1 hour. Playback rate: 5 frames per second (fps).

**Movie 3**. IGR1-RAC1^P29S^ melanoma cells (CellTrace-CFSE, green) co-cultured with heat-killed Jurkat T cells (CellTrace-Far Red, magenta) and brightfield image. Time is in minutes relative to start of the imaging (right after co-culture), and the time interval is 1.5 minutes. Scale bar: 20 microns. Playback rate: 20 frames per second (fps).

**Movie 4**. IGR1-RAC1^WT^ melanoma cells (CellTrace-CFSE, green) co-cultured with heat-killed Jurkat T cells (CellTrace-Far Red, magenta) and brightfield image. Time is in minutes relative to start of the imaging (right after co-culture). Scale bar: 20 microns. Playback rate: 20 frames per second (fps).

**Movie 5**. A magnified timelapse series of IGR1-RAC1^P29S^ melanoma cells (CellTrace-CFSE, green) co-cultured with heat-killed Jurkat T cells (CellTrace-Far Red, magenta) and brightfield image. Time is in minutes (min) relative to first frame shown. Scale bar: 20 microns. Movie shows an engulfment event where the phagocytic cup is visualized by inverted fluorescence (black, right) of CellTrace-CFSE. Playback rate: 20 frames per second (fps).

**Movie 6**. IGR1-RAC1^P29S^ melanoma cells (CellTrace-CFSE, green) co-cultured with living Jurkat T cells (CellTrace-Far Red, magenta) and brightfield image. Time is in minutes relative to start of the imaging (right after co-culture). Scale bar: 20 microns. Playback rate: 20 frames per second (fps).

**Movie 7**. IGR1-RAC1^WT^ melanoma cells (CellTrace-CFSE, green) co-cultured with living Jurkat T cells (CellTrace-FarRed, magenta) and brightfield image. Time is in minutes relative to start of the imaging (right after co-culture). Scale bar: 20 microns. Playback rate: 20 frames per second (fps).

**Movie 8**. A magnified timelapse series of IGR1-RAC1^P29S^ melanoma cells (CellTrace-CFSE, green) co-cultured with living Jurkat T cells (CellTrace-Far Red, magenta) and brightfield image showing a melanoma cell engulf a whole living Jurkat T cell. Time is in minutes (min) relative to the first frame shown. Scale bar: 20 microns. Playback rate: 20 frames per second (fps).

**Movie 9**. A magnified timelapse series of IGR1-RAC1^P29S^ melanoma cells (CellTrace-CFSE, green) co-cultured with living Jurkat T cells (CellTrace-Far Red, magenta) and brightfield image showing a melanoma cell engulf a piece of a Jurkat T cell. Time is in minutes (min) relative to the first frame shown. Scale bar: 20 microns. Playback rate: 20 frames per second (fps).

**Movie 10**. A magnified timelapse series of IGR1-RAC1^P29S^ melanoma cells (CellTrace-CFSE, green) co-cultured with living Jurkat T cells (CellTrace-Far Red, magenta) and brightfield image showing a T cell that dies as it interacts with the melanoma cell and then gets engulfed. Time is in minutes (min) relative to the first frame shown. Scale bar: 20 microns. Playback rate: 20 frames per second (fps).

**Movie 11**. A timelapse series with overlaying masks of IGR1-RAC1^P29S^ melanoma cells (CellTrace-CFSE, green) co-cultured with living Jurkat T cells (CellTrace-Far Red, magenta). White outlines mark masks of the green melanoma cells and free Jurkat T cells. Orange outlines mark the masks for T cells in contact or partially overlapping with melanoma cells. Cyan outlines marked masks that measure an engulfment event. Time is in minutes (min) relative to the start of the co-culture. Scale bar: 20 microns. Playback rate: 20 frames per second (fps).

**Movie 12**. A timelapse series with overlaying masks of IGR1-RAC1^WT^ melanoma cells (CellTrace-CFSE, green) co-cultured with living Jurkat T cells (CellTrace-Far Red, magenta). White outlines mark masks of the green melanoma cells and free Jurkat T cells. Orange outlines mark the masks for T cells in contact or partially overlapping with melanoma cells. Cyan outlines marked masks that measure an engulfment event. Time is in minutes (min) relative to the start of the co-culture. Scale bar: 20 microns. Playback rate: 20 frames per second (fps).**Figure 1. Dosing and sampling timeline**. Participants received doses at month 0 and month 2. Peripheral blood mononuclear cell samples for gE-specific CD4+ and serum samples for anti-gE antibody concentrations were collected on months 0, 2, 3, 14, 26, and 38. The study’s entire timeline lasted slightly more than three years.

